# An anatomical investigation of alkaptonuria: Novel insights into ochronosis of cartilage and bone

**DOI:** 10.1101/2024.09.11.612405

**Authors:** Juliette H Hughes, Gemma Charlesworth, Amanda Prior, Claire M Tierney, Paul D Rothwell, Neil P Thomas, Lakshminarayan R Ranganath, James A Gallagher, Alistair P Bond

## Abstract

Ochronotic pigmentation of connective tissue is the central pathological process in the rare metabolic disease alkaptonuria (AKU). Tissue pigmentation in AKU occurs due to unmetabolized homogentisic acid (HGA) in the circulation, caused by an enzyme deficiency in the liver. Ochronotic pigmentation, derived from HGA, has previously been reported and described in large joints obtained from arthroplasty surgeries, which typically have advanced disease. Many tissues that are affected by ochronosis are not accessible for study during life, including tissues subjected to early and mid-stage disease. Here, the opportunity arose to anatomically examine a 60-year-old AKU female body donor, allowing the investigation of previously understudied tissue, including those undergoing early-stage pathological changes. Dissection of fresh-frozen tissue was carried out and harvested tissues were fixed and examined histologically using H&E and Schmorl’s stains to aid identification of ochronotic pigment. This work focusses on osteochondral tissues including extra-skeletal cartilage, viscera and eyes. Gross and histological images demonstrating pigmentation in the cartilage and perichondrium of the ear ossicles, tympanic membrane, and the pubic symphysis fibrocartilaginous disc are described for the first time here. We also show the first examination of the temporomandibular joint, which macroscopically appeared unpigmented, with histological analysis of the fibrocartilaginous disc showing no pigmentation. Pigmentation of non-articular hyaline cartilage was observed in the respiratory tract, in both the hyaline cartilage and perichondrium, confirming previous findings. Within smaller joints, pigmentation of chondrons and the surrounding territorial matrix was observed, but was confined to calcified articular cartilage, and was not generally found in the hyaline articular cartilage. Dark pigmentation of the perichondrium adjacent to the articular surface was observed in numerous small joints, which has not been described before. The calcified bone matrix was not pigmented but ochronosis was identified in a small fraction of trabecular osteocytes in the capitate and radius, with substantially more pigmented osteocytes observed in bone of the ear ossicles. Viscera examined were unpigmented. This anatomical examination of tissues from an AKU individual highlights that most osteochondral tissues are susceptible to HGA-derived pigmentation, including the ear ossicles which are the smallest bones in the body. Within joints, calcified cartilage and perichondrium appear to be the earliest affected tissues, however why this is the case is not understood. Furthermore, why the TMJ disc was unaffected by pigmentation is intriguing. The heterogenous appearance of pigmentation both within and between different tissues indicates that factors other than tissue type (i.e. cartilage, perichondrium) and matrix composition (i.e. collagen-rich, calcified) may affect the process of ochronosis, such as oxygen tension, loading patterns and tissue turnover. The effect of nitisinone treatment on the ochronotic disease state is considered, in this case 7 years of treatment, however comparisons could not be made to other cases due to inter-individual variability.

## 2 Introduction

Alkaptonuria (AKU; OMIM #203500) is an ultra-rare metabolic disease that results in degeneration and failure of connective tissues (Phornphutkul et al., 2002). Autosomal recessive mutations in the homogentisate 1,2-dioxygenase (HGD) gene lead to an inability to breakdown homogentisic acid (HGA) in AKU (La Du et al., 1958, Fernández-Cañón et al., 1996, Zatkova et al., 2012). Unmetabolized HGA is excreted into the urine, but it still elevated in the circulation and tissue fluid. The deposition of a yellow (or “ochre”) to dark brown/black pigment into connective tissues, a process called ochronosis, is characteristic of AKU and ultimately causes devastating joint destruction and connective tissue disorder (Virchow, 1866, O’Brien et al., 1963) and heart valve disease (Phornphutkul et al., 2002, Hannoush et al., 2012, Helliwell et al., 2008). In early adulthood, backpain and joint pain begin as early as the third decade (Cox et al., 2019), progressing to a severe and early-onset osteoarthropathy of the spine and large joints, resulting in joint replacement and immobility of the spine (Phornphutkul et al., 2002, Mannoni et al., 2004). Other affected tissues include tendons and ligaments leading to rupture, and the heart valves causing stenosis. Collectively these pathological changes contribute to deteriorating quality of life.

Ochronosis of connective tissue, such as the sclera of the eye and cartilage of the ear, typically takes two to three decades to become visible externally, with ear cartilage biopsies from adult AKU patients detecting pigmentation at the age of 20 years (Cox et al., 2019). It is however noteworthy that a recent paediatric study reported scleral pigmentation in 3 out of 13 AKU children, with the earliest observed at 13 years, with no externally visible pigmentation of ear cartilage (Kujawa et al., 2023). Due to the location of articular cartilage and other joint tissues, they cannot easily be studied, particularly at the microscopic level, to determine when pigment is first deposited. Despite the rarity of AKU, the ochronotic phenotype of AKU joints donated after surgery has been documented to some degree, offering valuable insights into the disease process within articular cartilage (Taylor et al., 2011, Boyde et al., 2014, Taylor et al., 2017, Taylor et al., 2019, Chow et al., 2020). Other tissues however are either not replaceable or do not cause severe enough symptoms to warrant replacement and are therefore unavailable to study (i.e. spine, respiratory cartilage). Other than invasive medical examinations, such as arthroscopy and biopsy, which require medical reasons to carry out such procedures, surgical insights and post-mortems are the only other opportunities to investigate ochronosis and related pathologies.

Many surgical case reports exist in the literature, often coupled with a brief history of the patient and radiographical findings and symptoms. Examples include bronchoscopy (Parambil et al., 2005), meniscus rupture and arthroscopy (Nag et al., 2013, Xu et al., 2015), arthroplasty (Merolla et al., 2012, Cebesoy et al., 2014, Khalifa et al., 2022) and tendon/ligament rupture (Manoj Kumar and Rajasekaran, 2003, Jiang et al., 2019, Mwafi et al., 2021). Such case reports vary in length and detail, with some showing gross photographs, with even fewer showing histology. Detailed descriptions of AKU patients both ante-mortem and post-mortem published approximately 70 years ago document individuals with severe disease but lack both high quality gross photographs and clear, high resolution histology photomicrographs (Galdston et al., 1952, Lichtenstein and Kaplan, 1953). A more recent post-mortem examination of an AKU patient was carried out approximately 15 years ago, providing the first modern gross and histological insight into the appearance of tissue ochronosis (Helliwell et al., 2008). Overall, the available literature provides a good summary of the location of ochronotic pigment across different body tissues at the macroscopic level, however more investigation into tissue ochronosis is needed to characterise the microscopic location of ochronotic pigment to gain further insight into the disease process.

Nitisinone (Orfadin®) has recently been approved for treatment of AKU (European Medicines Agency, 2020). It inhibits the 4-hydroxyphenylpyruvate dioxygenase (HPPD) enzyme upstream of HGD, preventing the conversion of 4-hydroxyphenylpyruvic acid (HPPA) into HGA (Ranganath et al., 2014, Ranganath et al., 2020c), thus decreasing HGA in urine and importantly the serum. A pre-clinical study demonstrated that nitisinone treatment of AKU mice was able to either prevent joint ochronosis if treated from birth or prevent further pigment deposition if treated midlife (Keenan et al., 2015, Preston et al., 2014). Although nitisinone is not approved for use in children, for younger adult patients with early-stage connective tissue ochronosis, nitisinone treatment will halt further pigment deposition and may prevent future connective tissue damage. For patients already living with significant ochronotic disease, connective tissue disorder and damage, such as osteoarthritis, and is unlikely to repair, therefore deterioration of tissue is likely to be slowed rather than halted by nitisinone. It has been shown that nitisinone slows the rate of disease progression in AKU and may increase quality of life and physical function, however it has not been able to halt or reverse disease progression in already affected patients (Ranganath et al., 2020b, Ranganath et al., 2020c, Spears et al., 2024). Currently, connective tissue disorder in AKU can only be treated by palliative strategies such as pain management, total joint arthroplasty, ligament and tendon repair after rupture, and heart valve replacement (Ranganath et al., 2013). None of these treatments intervene in the pathophysiological process of ochronosis and subsequent biomechanical failure of tissues, but intervene at the point of, or after, failure of the tissue.

Although it is known that AKU patients have accelerated ageing and degeneration of joints and other connective tissues, the process of ochronosis is not well understood, and neither are the changes that lead to tissue dysfunction. It is not known where in the extracellular matrix HGA-derived pigment binds, nor has the chemical composition of pigmentation been identified (Ranganath et al., 2019). It is clear however that AKU tissue ages and degenerates rapidly, even quicker than the general osteoarthritic population. Understanding AKU pathophysiology may be the key towards development of therapies that prevent tissue degeneration in AKU, that also may be applicable to other general, degenerative diseases such as osteoarthritis, tendinopathy and valve stenosis, where common mechanisms may exist that are amplified by AKU. Even though a surprising number of papers/case reports over many years have documented pigmented tissues in AKU, most have only been able to grossly describe the location of pigmentation and the clinical signs of tissue pathology that result. Few have examined the tissue using histology or other methods to aid understanding of the pathophysiology. There are also many tissues that cannot be studied, because they are not replaceable or are not considered severely diseased enough to warrant surgical repair.

Here, we were provided with an exceptionally rare opportunity to examine tissues from an AKU body donor. Anatomical and histological examination of the organs and tissues was carried out, to describe the distribution of ochronotic pigmentation and related pathology with a focus on osteochondral tissue, including previously unexamined tissues.

## 3 Materials and methods

### 3.1 Ethical approval

A 60-year-old female with alkaptonuria donated her body post-mortem to the University of Liverpool for anatomical examination. Ethical approval was obtained from the University of Liverpool Central Research Ethics Committee (reference number: 5785). Consent was obtained prior to death, which included the use of photographs for research purposes.

### 3.2 Donor information

The individual had been receiving nitisinone treatment (2mg/daily) for 7 years. Prior to nitisinone treatment, serum HGA was 42.6 µmol/L and 24-hour urinary HGA was 24.2 mmol/L. Following 72 months of nitisinone treatment (2mg/daily) serum HGA decreased to <3.1 µmol/L and 24-hour urinary HGA was 0.4 mmol/L.

### 3.3 Dissection

The donor was refrigerated for 7 days prior to arriving at University of Liverpool, where upon the body was placed in −20°C for storage. The body was defrosted in the refrigerator, prior to dissection, which was carried out over 5 days, in a regional manner. Photographs were taken of anatomical tissues and structures *in situ* and after removal. Smaller samples were removed for histological analysis.

### 3.4 Histology

Samples from dissected tissues were fixed in 10% phosphate buffered formalin for a minimum of 48 hours, and subsequently stored in 70% ethanol until processing. Tissues that contained mineral were decalcified prior to tissue processing. All tissues were decalcified by immersion in Formical-2000 (Statlab, US) until mineral was no longer present, which varied from several hours to several days with replenishment of the solution depending on the size of the tissue. Tissues were processed using a Leica ASP300 tissue processor. Briefly, tissues were dehydrated by immersion in 70% - 100% ethanol, immersed in xylene and then transferred to molten Formula R paraffin wax (Leica, Germany). All viscera were processed using protocol 1, and a longer protocol was used for bone and cartilage (protocol 2) and a separate protocol for the sclera (protocol 3), see Supplementary Tables 1 and 2 for details. Tissues were embedded in Formula R paraffin wax (Leica, Germany) and sectioned at 4.5 – 5 µm using a Leica RM2235 microtome and mounted on glass Superfrost® Plus microscope slides (ThermoScientific, UK).

Sections were stained for ochronotic pigment using a modified Schmorl’s stain, which stains the yellow-brown pigment a dark blue-green colour (Tinti et al., 2011, Hughes et al., 2021). Briefly, sections were deparaffinised, rehydrated, incubated in Schmorl’s stain (1% ferric chloride, 1% potassium ferricyanide in distilled water), immersed in 1% acetic acid and counterstained with nuclear fast red (full protocol in supplementary information). Sections were stained with haematoxylin and eosin (H&E) following deparaffination and rehydration, see supplementary information for full protocol. Slides were imaged using a Zeiss Axioscan Z1 slide scanner and processed using Zen 3.8 software (Zeiss).

## 4 Results

Pigmentation of tissue typically appears as a black/brown colour grossly, and when sectioned, appears as a golden yellow or brown colour. H&E staining can mask less intense pigment. Schmorl’s staining was therefore carried out alongside H&E, to stain the pigment a green-blue colour, aiding its identification (Tinti et al., 2011, Taylor et al., 2012, Preston et al., 2014, Hughes et al., 2021).

### 4.1 Temporomandibular joint

Upon gross examination of the temporomandibular joint (TMJ), a synovial joint of the head and neck, the articular surfaces of the mandibular fossa/articular tubercle of the temporal bone and the mandibular head of the mandible did not appear pigmented (not shown). The articular surfaces of the TMJ were not examined histologically. The articular disc of the TMJ, which is composed of a dense fibrocartilaginous-like fibrous connective tissue (Runci Anastasi et al., 2021, Detamore and Athanasiou, 2003), appeared normal, both grossly and histologically, with no pigmentation identified with Schmorl’s staining (Supplementary Figure 1). Note that fibrocartilage does not have a perichondrium.

### 4.2 Ear ossicles

The tympanic membrane attached to the malleus was pigmented both grossly and histologically, particularly at its attachment to the handle of the malleus at the umbo and lateral process (Figure 1a-f), and also towards the periphery of the membrane. Pigmentation in places appeared to be orientated with fibre direction (Figure 1f). Figure 1d-e were annotated with reference to the histological images by De Greef et al. (2016) and Graham et al. (1978), who investigated the connection between the tympanic membrane and malleus, termed the tympano-mallear connection. The malleus was very pigmented where the tympanic membrane attaches (Figure 1d-e), with dark pigmentation in the perichondrium lining the cartilage that encases the bone forming the distal end of the malleus handle. Chondrocyte pigmentation was observed. Dense regular connective tissue (DCT) within the tympanic membrane could be observed, particularly with H&E staining (Figure 1e), in addition to loose connective tissue (LCT) pigmentation, that is not as eosinophilic as DCT. In the H&E stained tympano-mallear connection, the DCT can be seen blending with the perichondrium. The DCT of the tympanic membrane was pigmented, whilst the LCT and the simple squamous epithelium was not. The articular facet of the malleus that articulates at the synovial incudo-malleolar joint was grossly unpigmented (Figure 1a), however pigmented chondrons were observed histologically within the calcified articular cartilage of the facet (Figure 1g). Most of these pigmented chondrons were deep to the tidemark, with a few pigmented chondrons observed superficially within the hyaline articular cartilage (HAC).

**Figure 1.**
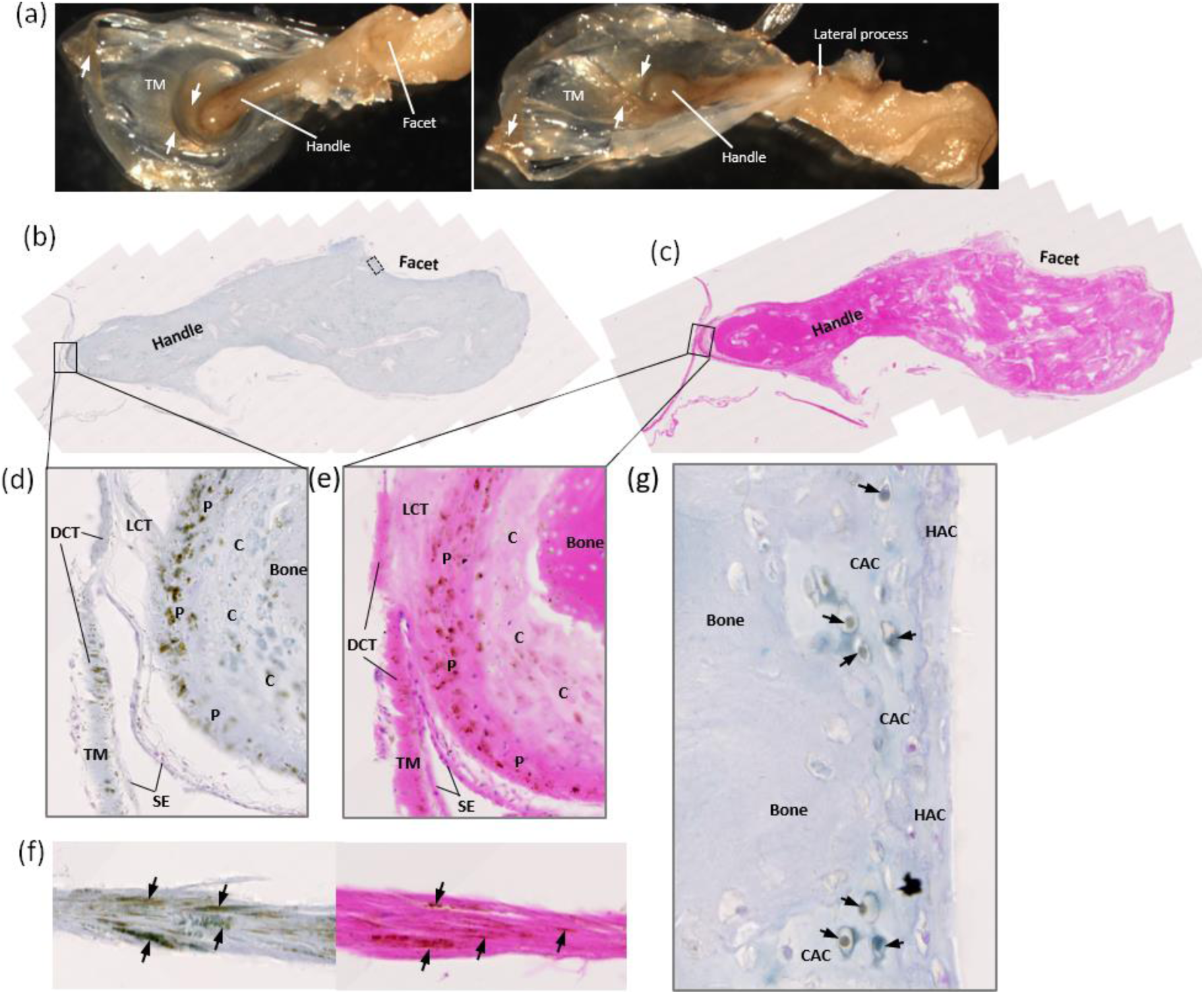
Ochronotic pigmentation of the malleus and tympanic membrane. The malleus is shown in (a) with the attached tympanic membrane (TM), which appears pigmented (white arrows) from the central attachment of the handle of the malleus, radiating outwards. The tympanic membrane also appears pigmented at the periphery. The lateral process of the malleus can be observed in the right image, and also appears pigmented. (b) shows Schmorl’s staining and (c) shows H&E staining of the whole malleus sectioned longitudinally, including part of the TM at its attachment to the handle of the malleus and the facet for articulation with the incus. (d) and (e) show higher magnification images of the attachment of the TM at the umbo/tip of the handle of the malleus. The TM is composed of a dense connective tissue (DCT) that appears pigmented. The tip of the handle of the malleus has been labelled to show the bone, cartilage (C), the perichondrium (P), with intense pigmentation observed in the perichondrium and the cartilage. Intervening between the DCT and perichondrium is loose connective tissue (LCT), which appears unpigmented. Lining both the TM and perichondrium is a layer of unpigmented, simple squamous epithelium (SE). (f) shows Schmorl’s (left) and H&E (right) staining of the TM near to the lateral process of the malleus, where intense pigmentation is observed running in the direction of fibre orientation of the TM. (g) shows pigmented chondrons (black arrows) within the calcified articular cartilage (CAC) of the articular facet of the malleus from the inset area shown in (b), with no pigmented chondrons observed in the hyaline articular cartilage (HAC).

Grossly, both the incus (Figure 2a) and stapes (Figure 2h) appeared normal. Histologically, pigmentation of the articular facet of the incus that articulates with the malleus was observed (Figure 2b-g), with pigmented chondrons observed in the CAC layer, and not the superficial HAC. The lenticular process of the incus was not examined histologically. The head of the stapes, that forms an articulation with the long limb of the incus at the incudostapedial joint, and the footplate (or base) of the stapes that rests on the oval window, can be observed in Figure 2h and histologically in Figure 2i-k, where pigmentation of chondrons within the CAC can be observed. In the footplate of the stapes, pigment was confined to CAC, and not present in the HAC or bone. In the head of the stapes, pigmentation surrounding a chondrocyte in HAC was observed.

**Figure 2.**
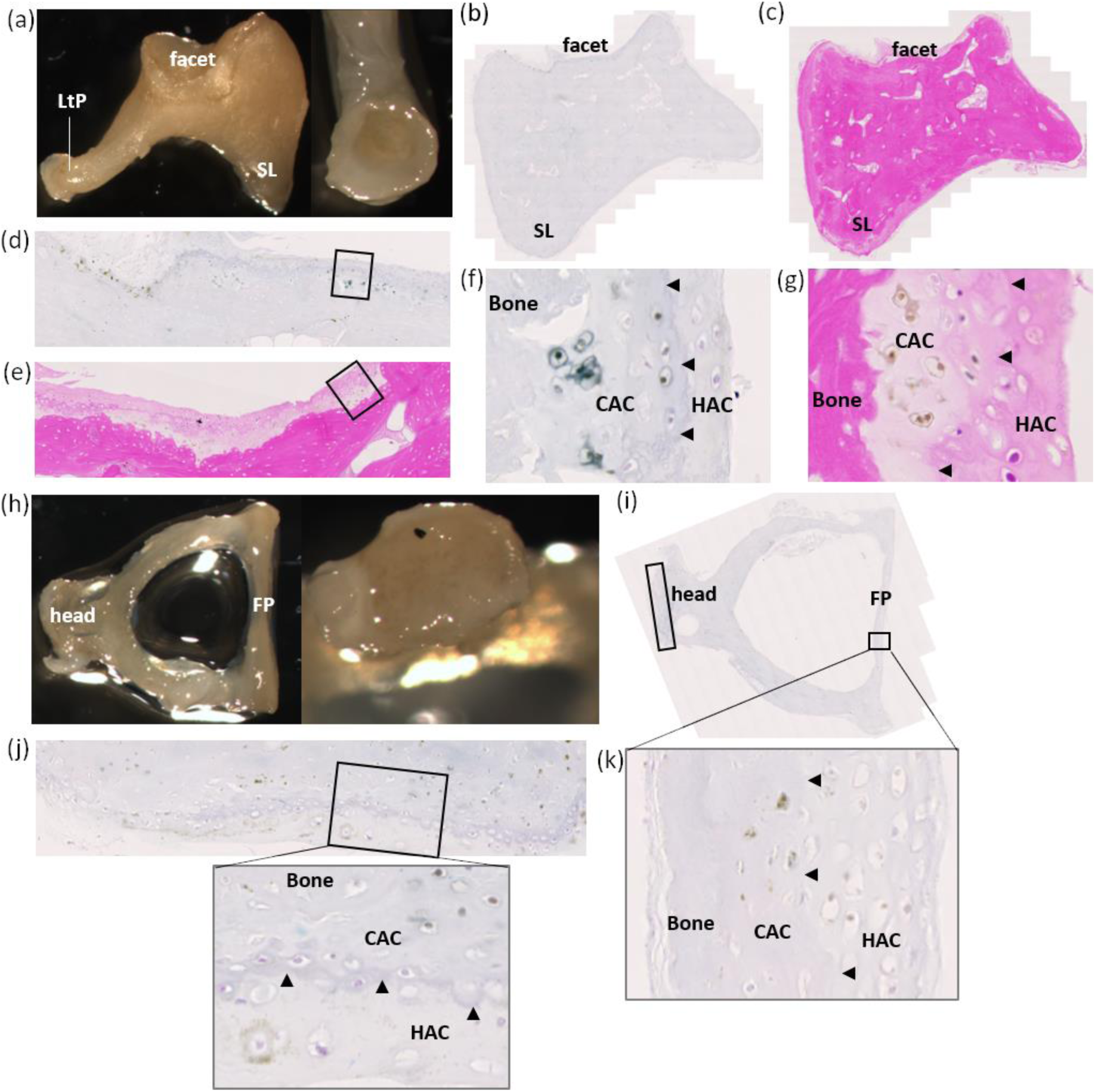
Ochronotic pigmentation of the incus and stapes. The incus is shown in (a), where the short limb (SL), lenticular process (LtP) and facet can be observed, with the right image showing the articular surface of the lenticular process (LtP). Macroscopically, no pigmentation was identified in the incus. (b) and (c) show H&E and Schmorl’s staining of the incus, with the articular facet for the malleus towards the top of the image and the short limb identified (SL). (d) and (e) show the articular facet of the incus stained with Schmorl’s and H&E, respectively, with the insets shown at higher magnification in (f) and (g). Pigmentation of chondrocytes within calcified articular cartilage (CAC) was observed, with no pigmentation observed in the hyaline articular cartilage (HAC), superficial to the tidemark (indicated by arrow heads). The stapes is shown in (h), with the head and footplate annotated, with the right image showing the distal end of the head of the stapes which appears unpigmented. (i) shows Schmorl’s staining of the stapes, with higher magnification images of the insets shown in (j) and (k). (j) shows the head of the stapes, which articulates with the incus. Pigment can be observed in the CAC, and also in the HAC superficial to the tidemark (indicated by arrowheads), surrounding a chondrocyte. (k) shows the footplate of the stapes, which shows pigmentation of chondrons in the CAC (tidemark indicated with arrowheads).

When examining the ear ossicles, it was also observed that osteocytes within the bone were pigmented. Figure 3a-c shows an example of pigmented osteocytes in the malleus, incus and stapes, with both H&E staining and Schmorl’s staining showing pigmented cells within the bone matrix (see arrows). Whilst many pigmented osteocytes were observed, there were still many non-pigmented osteocytes.

**Figure 3.**
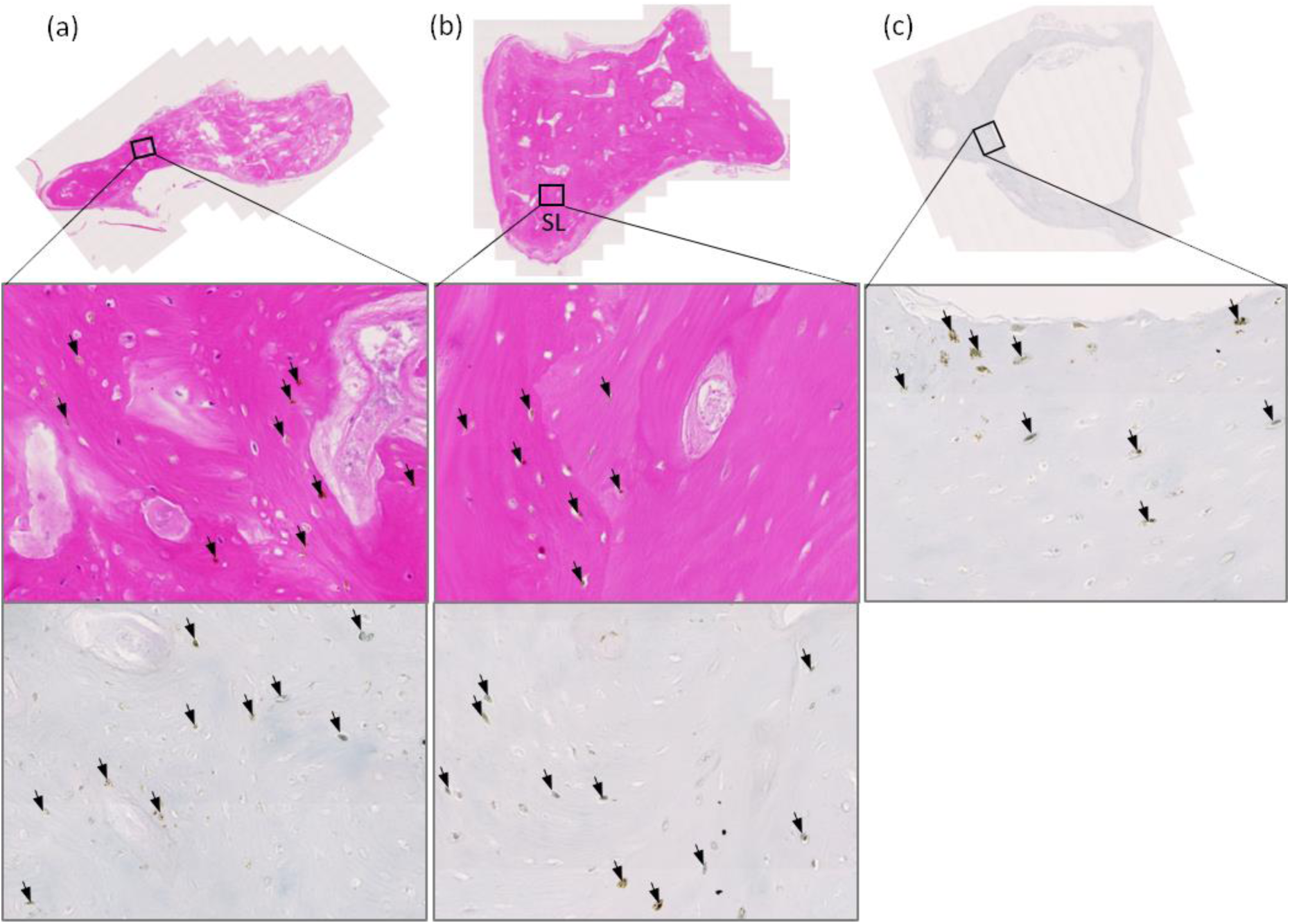
Ochronotic pigmentation of ear ossicle osteocytes. The malleus and incus stained with H&E in (a) and (b), respectively. The insets in (a) and (b) are shown below, with a corresponding area stained with Schmorl’s. The stapes stained with Schmorl’s in (c), with the inset shown below. Arrows indicate pigmented cells (osteocytes) within the bone, which can be observed with both H&E and Schmorl’s staining.

### 4.3 Pubic symphysis

In a coronal cross-section through the pubic symphysis (Supplementary Figure 2), the joint looked unpigmented, including the fibrocartilaginous disc. The superior part of the pubic symphysis examined histologically showed pigmentation within the fibrocartilaginous disc that was intense and patchy, and associated with cells, assumed to be chondrocytes (Supplementary Figure 2b-c). The disc had a cleft, which is considered a normal feature in some individuals (Becker et al., 2010). The thin hyaline cartilage layer showed some very light pigmentation of chondrocytes with Schmorl’s staining (Supplementary Figure 2d), however very few cells showed pigmentation. The bone appeared normal.

### 4.4 Costal cartilage

All costal cartilage observed was intensely pigmented and black in colour, both at the costal margin and sternocostal junctions (Figure 4a-b), and was also very hard and brittle. An unstained section of the black cartilage of the left second sternocostal junction shown in Figure 4c, histologically shows golden yellow or ochre coloured pigment, spread throughout the entire cartilage matrix (Figure 4d). H&E and Schmorl’s staining of the same area confirmed that chondrocytes and the entire HAC matrix was pigmented (Figure 4e-f). A clear delineation between the outer and inner layers of the thick perichondrium present around costal hyaline cartilage was observed (Figure 4e-f). The inner chondrogenic layer showed pigmented chondroblasts and pigmented extracellular matrix. The outer fibrous perichondrium showed pigmented fibroblasts and little pigmentation within the extracellular matrix.

**Figure 4.**
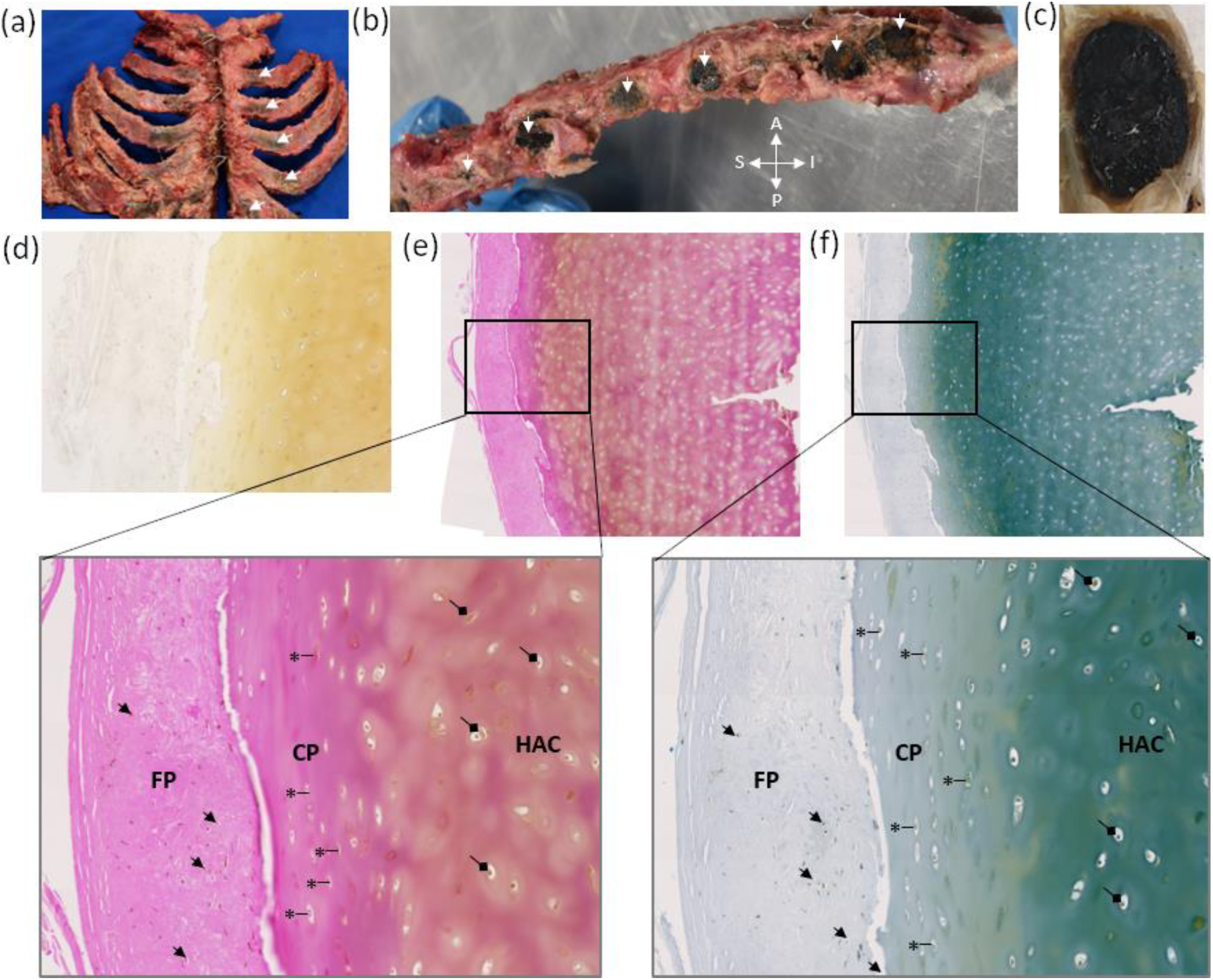
Intense ochronotic pigmentation of costal cartilage. An anterior view of the sternum and associated costal cartilage and ribs is shown in (a), with intense black pigmentation of costal cartilage indicated by arrows on the left side, with pigment also present on the right side. A lateral view of the sternum with the ribs removed at the sternocostal junctions is shown in (b), with arrows indicating black pigmented cartilage (A = anterior, P = posterior, S = superior, I = inferior). The costal cartilage of left second rib was transected and is shown in (c) with intense black pigmentation observed throughout the cartilage. Histological sectioning of the left second costal cartilage in (c) is shown in (d) with no staining where pigment appears a yellow-ochre colour, in (e) with H&E staining and (f) with Schmorl’s staining, with the insets shown at greater magnification below. Intense pigmentation can be observed throughout the hyaline articular cartilage (HAC) of the costal cartilage, within chondrocytes (indicated by diamond arrows) and spread through the entire matrix. Surrounding the HAC is a thick perichondrium, divided into two layers; an outer fibrous perichondrium (FP) where pigmentation is mostly associated with fibroblast cells (examples indicated by arrows) and only a small amount in the matrix, and an inner chondrogenic perichondrium (CP), where pigmentation is both within chondroblasts (examples indicated by an asterix) and throughout the matrix.

### 4.5 Elastic cartilage

The auricles and lateral part of the external auditory meatus are composed of elastic cartilage. Grossly, the right ear overall was much darker than the left ear, with the cartilage also appearing darker in colour when a circular biopsy from the conchal bowl of each ear was bisected (Figure 5a-b). A thin slice from each conchal biopsy was examined via dark field illumination using a dissection microscope (Figure 5c-d), with a direct comparison shown in Figure 5e. The cartilage pigmentation was patchy and unevenly distributed, and more intense in the right biopsy, which was confirmed with Schmorl’s staining (Figure 5f-g), with an area of more intense pigmentation highlighted by arrows in Figure 5f. Small areas of the perichondrium were pigmented in both biopsies, see the arrow heads in Figure 5g. The skin adjacent to the cartilage appeared unpigmented in both biopsies. Grossly, pigmentation was only observed at the periphery of the elastic cartilage of the external acoustic meatus in the perichondrium (Figure 6a), also shown histologically in Figure 6b-c, with the elastic cartilage itself appearing normal. Note that some of the perichondrium remained unpigmented. Histologically, no pigmentation was identified within the external acoustic meatus cartilage chondrocytes or extracellular matrix.

**Figure 5.**
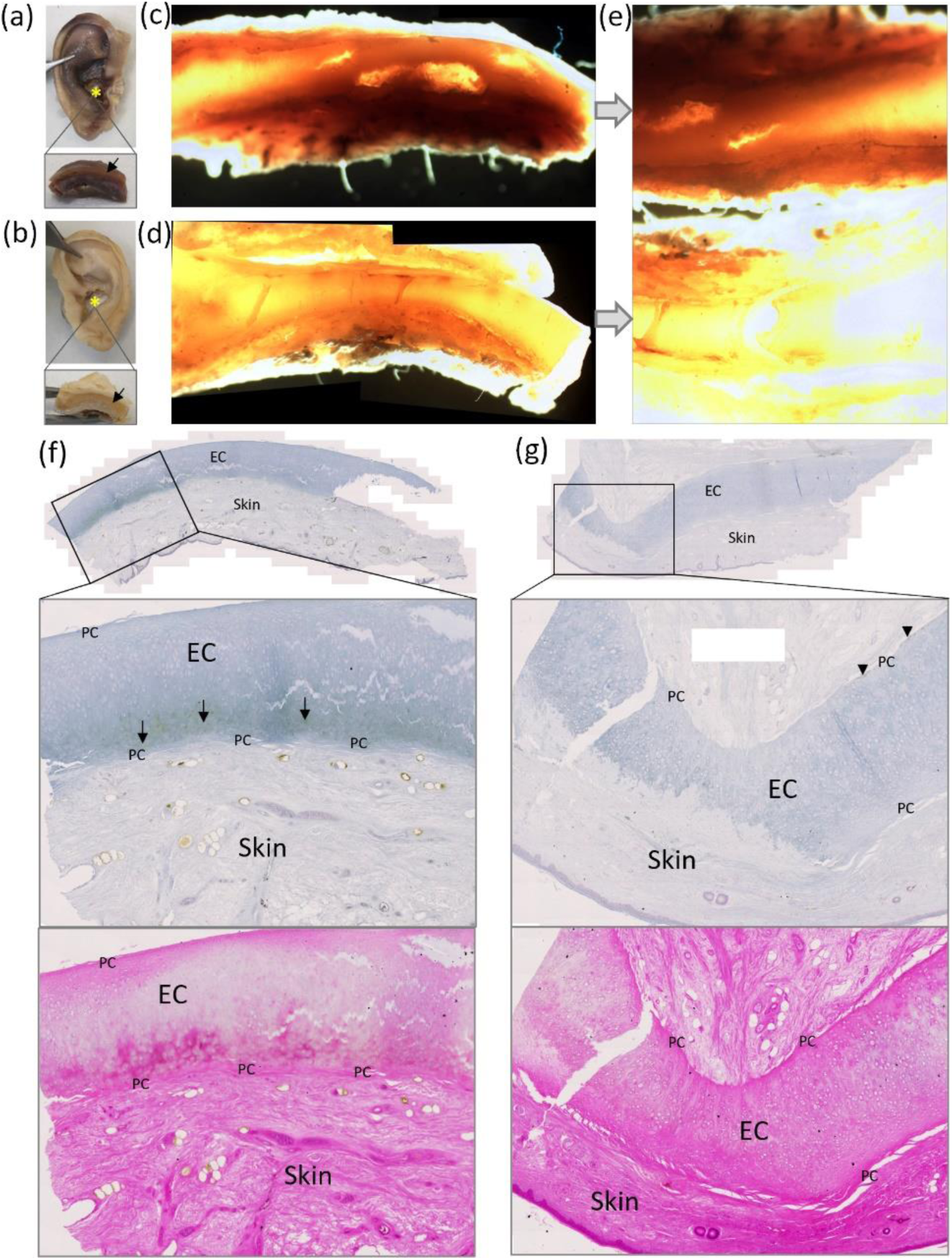
Ochronotic pigmentation of elastic cartilage from the ear. The right and left intact auricles are shown in (a) and (b) respectively (post-fixation). The asterix indicates where a circular biopsy approximately 1cm in size was removed from the conchal bowl of each auricle, and then bisected, with the cross section shown in the inset and the arrow indicating the elastic cartilage. A thin slice of each biopsy was examined with a dissecting microscope using dark field illumination and photographed; right shown in (c) and left shown in (d), with the right appearing much darker suggesting more pigmentation. A direct comparison of the right (top) and left (bottom) biopsies is shown in (e). Histological sections of the right and left conchal bowl biopsies, stained with Schmorl’s stain, are shown in (f) and (g) respectively, with the inset shown at greater magnification below with a corresponding H&E stained image of the same area. Arrows in (f) demonstrate the most pigmented area of the elastic cartilage (EC), adjacent to perichondrium (PC). Arrow heads in (g) indicate mild pigmentation of perichondrium.

**Figure 6.**
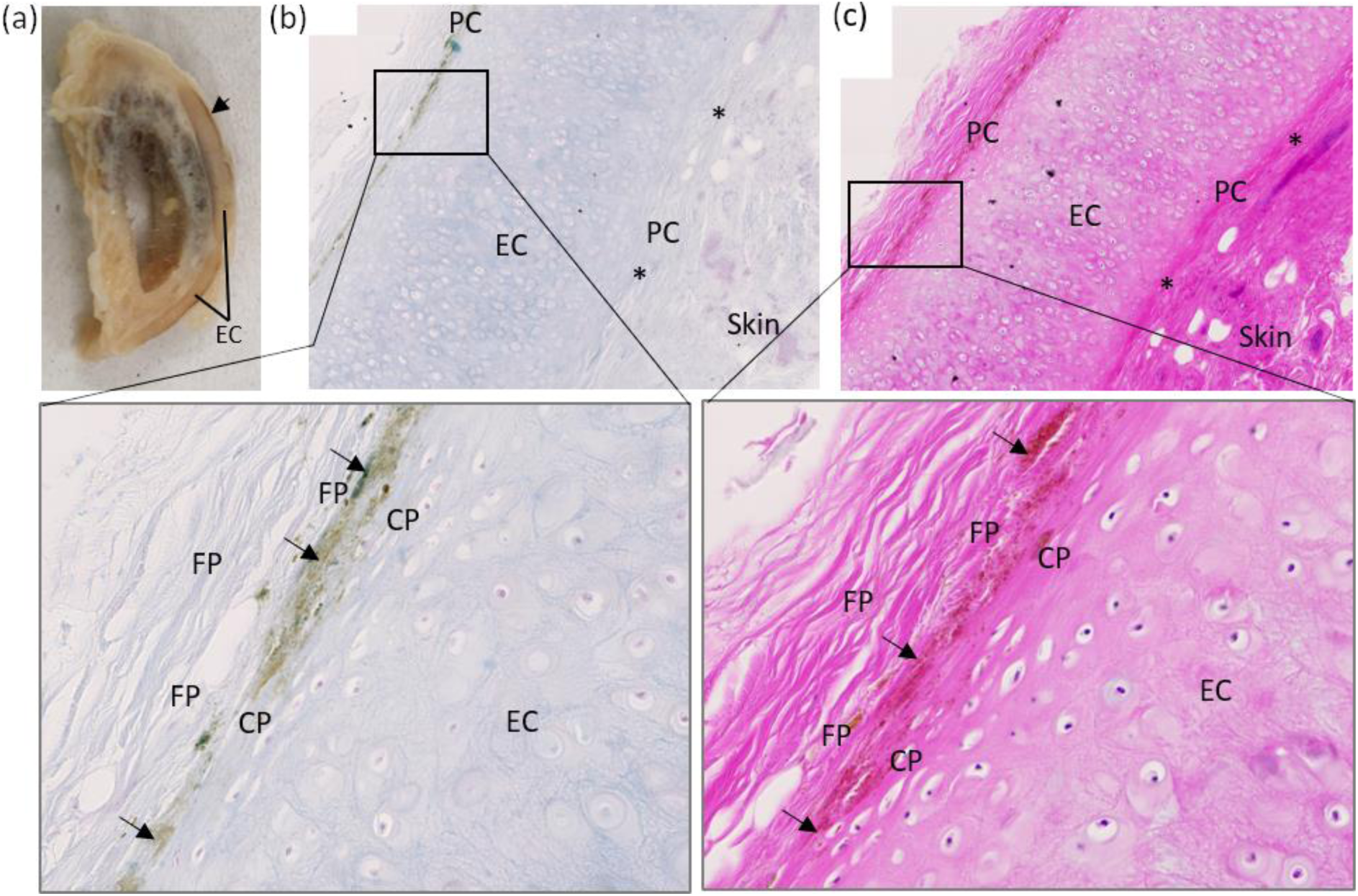
Ochronotic pigmentation of elastic cartilage perichondrium. A sagittal section through the right external acoustic meatus is shown in (a), with the arrow highlighting an area of pigmentation observed macroscopically at the periphery of the elastic cartilage (EC). Schmorl’s staining of the external acoustic meatus elastic cartilage (EC) is shown in (b) with a corresponding area stained with H&E in (c). Pigmentation only appears to be associated with the perichondrium, situated mostly within the outer fibrous perichondrium (FP) adjacent to the boundary with the inner chondrogenic perichondrium (CP), although some appears to be located within the chondrogenic perichondrium. The asterix indicates areas of perichondrium that are not pigmented.

### 4.6 Respiratory cartilage

Hyaline cartilage of the respiratory tract was examined. Dark pigmentation at the periphery of the nasal septum cartilage was observed grossly, with histological analysis showing no pigmentation associated with the chondrocytes or the cartilage matrix (Figure 7a). Grossly, the thyroid cartilage of the larynx and adjacent upper tracheal rings were intensely pigmented and were black in colour (Figure 7b). Histologically, cartilage pigmentation of a superior mediastinum tracheal ring was identified (Figure 7b). Pigmentation was observed in cartilage from the primary bronchus of the left bronchial tree, but was not identified in secondary bronchus cartilage (Figure 7c-d). Perichondrium pigmentation was observed in all of the respiratory cartilages examined histologically (nasal septum, tracheal ring, bronchial cartilage), situated between the inner chondrogenic and outer fibrous layers (Figure 7a-d).

**Figure 7.**
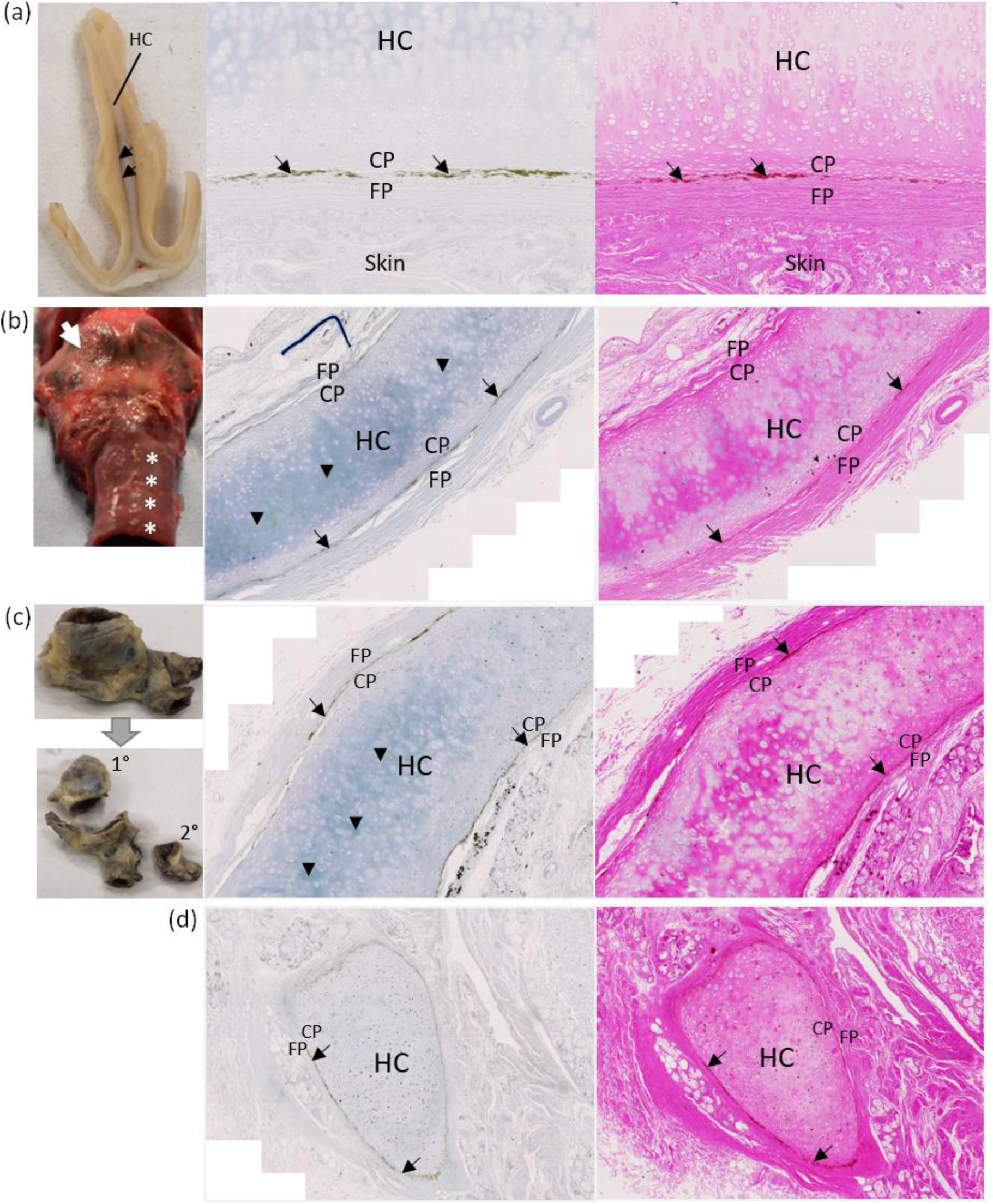
Ochronotic pigmentation of hyaline respiratory cartilage and perichondrium. Gross images of hyaline cartilage from the respiratory tract are shown on the left of each row, with corresponding Schmorl’s (middle) and H&E (right) staining. A coronal section through the nasal septum (post-fixation) is shown in (a) with the location of hyaline cartilage (HC) shown and arrows indicating pigmentation at the periphery of the HC. Histologically, no pigmentation was observed within the HC, but was observed within the perichondrium, situated at the boundary between the fibrous perichondrium (FP) and the chondrogenic perichondrium (CP). The larynx and proximal trachea (pre-fixation) are shown in (b), with the arrow indicating the thyroid cartilage and the asterix’s indicating the most superior tracheal rings, all of which appear black with pigmentation. The histology in (b) shows a tracheal ring from the superior mediastinum, with pigmentation observed within the hyaline cartilage (HC) indicated by arrow heads, and arrows indicating pigmentation of the perichondrium located between the FP and CP. A segment of the left bronchial tree, with primary (1°) and secondary (2°) bronchi isolated, is shown in (c), although the cartilage cannot be observed grossly from this view. Pigmentation was observed within the primary bronchial cartilage as indicated by arrows heads in row (c) via Schmorl’s staining, but not within secondary bronchial cartilage in row (d). Pigmentation was observed in the perichondrium between the FP and CP in both primary and secondary bronchial cartilage.

### 4.7 Upper limb joints

The shoulder joints were both artificial and were therefore not examined. Examination of the left upper limb showed pigmentation of articular cartilage of the distal humerus, radial head and proximal ulna as golden brown to dark brown pigment (Figure 8a-b). Overall, pigmentation was confined mostly to the edges of the articular surfaces of upper limb joints, see Figure 8a,c-d, although in the radial head, dark pigmentation was observed on the main articular surface, see arrows in Figure 8b. Most of the articular cartilage of the capitate grossly appeared unpigmented, however when examined histologically pigmentation of chondrons was observed in the calcified articular cartilage, with both intracellular, pericellular and territorial matrix pigmentation (Supplementary Figure 8). Histological examination of the distal 5^th^ interphalangeal joint revealed pigmented chondrons within the articular cartilage (data not shown).

**Figure 8.**
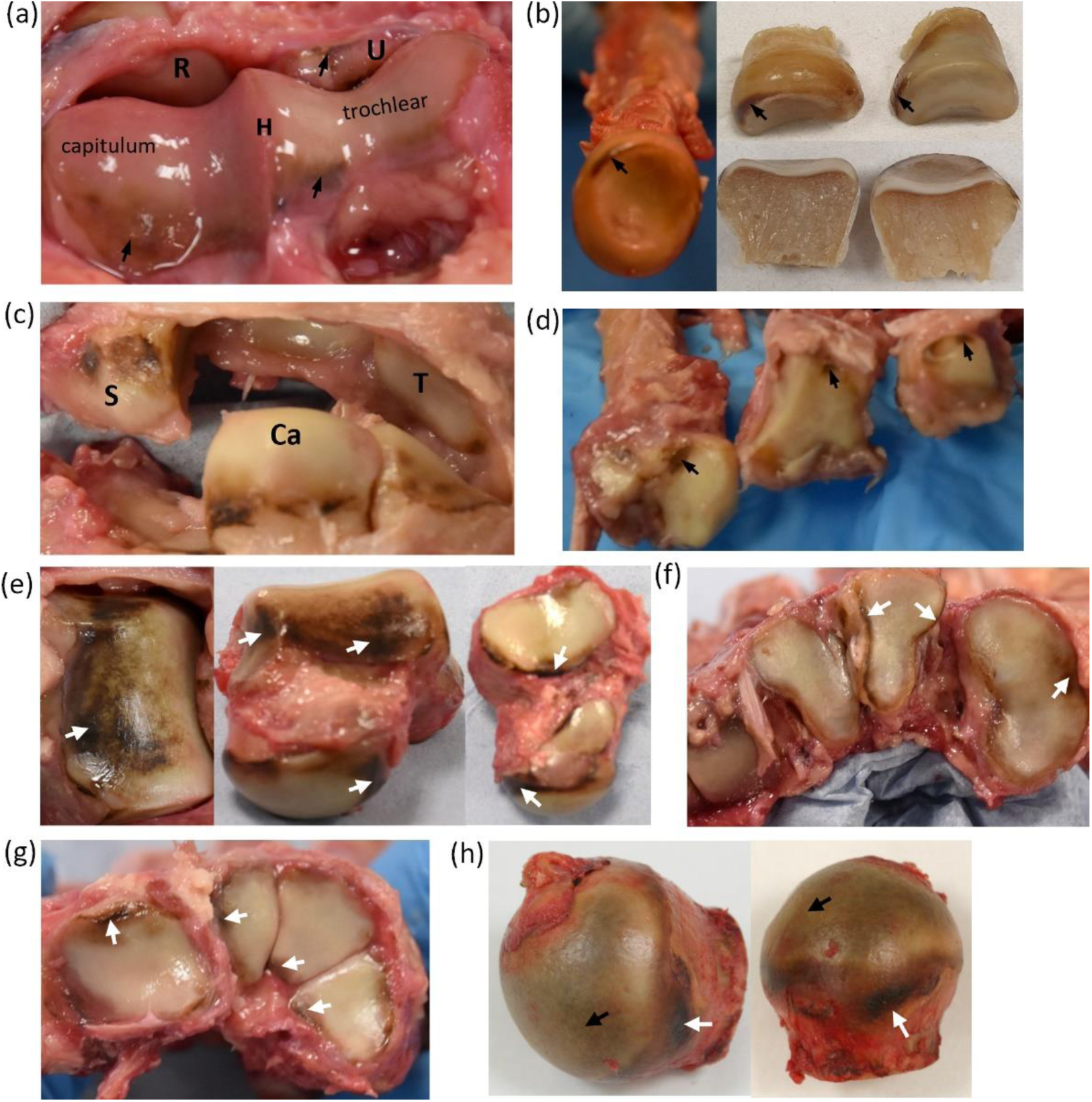
Ochronotic pigmentation of upper and lower limb articular cartilage and perichondrium. A superior view of the distal end if the left humerus (H) is shown in (a), with pigmentation indicated by arrows on the articular cartilage of the humerus. The proximal ends of the radius (R) and ulna (U) are visible. A superior view of the radial head is shown in (b) (left), which has been bisected (right), with pigmentation of articular cartilage indicated by arrows. The scaphoid (S), capitate (Ca) and triquetral (T) carpal bones are shown in (c), with arrows indicating dark pigmentation that is located at the periphery of the articular surface, adjacent to the bone. The proximal ends of the 2nd to 4th metacarpal bones, from left to right, are shown in (d), with pigmentation present at the edge of the articular surfaces, indicated by arrows. From left to right, (e) shows superior, anterior and inferior views of the talus with arrows indicating areas of pigmentation. The proximal ends of the 1st to 4th metatarsals (from right to left) are shown in (f), with arrows indicating pigmentation. The proximal ends of the cuboid, medial cuneiform, intermediate cuneiform and lateral cuneiform tarsal bones (from left to right) are shown in (g). The femoral head of the hip joint is shown in (h), with black arrows highlighting pigmentation of the main articular surface, and the white arrows indicating pigmentation of perichondrium. All gross images were taken pre-fixation, except the bisected radial head images taken post-fixation in (b).

When bisected and photographed, macroscopic pigmentation could be observed deep within the radial head articular cartilage, immediately adjacent to the unpigmented subchondral bone (Figure 9a). Figure 9b-e shows pigmentation associated with chondrocytes in the CAC, and not within the HAC superficial to the tidemark. Pigmentation is more advanced in some chondrons than others, with only pericellular pigmentation observed in some chondrons compared to pigment spreading into the territorial and inter-territorial matrix of other chondrons. Pigmented chondrons were observed adjacent to the cement line, with the most heavily pigmented chondrons in Figure 9b appearing to be in the deepest areas of the CAC, furthest from the tidemark; this was also observed in the capitate histology (Supplementary Figure 3a). The tidemark was also duplicated, and in some regions there were 3 tidemarks (not shown), with a few lightly pigmented chondrons observed between the tidemarks (Figure 9, Supplementary Figure 3). In Figure 9c, heavily pigmented chondrons were observed (see the inset), with the pigment remaining unstained with Schmorl’s stained and appearing dark gold-brown in colour. A noteworthy feature was also identified, with a line of pigmentation appearing to bridge vertically between two pigmented chondrons and from the most superficial chondron to the tidemark, see the black arrows in inset of Figure 9c.

**Figure 9.**
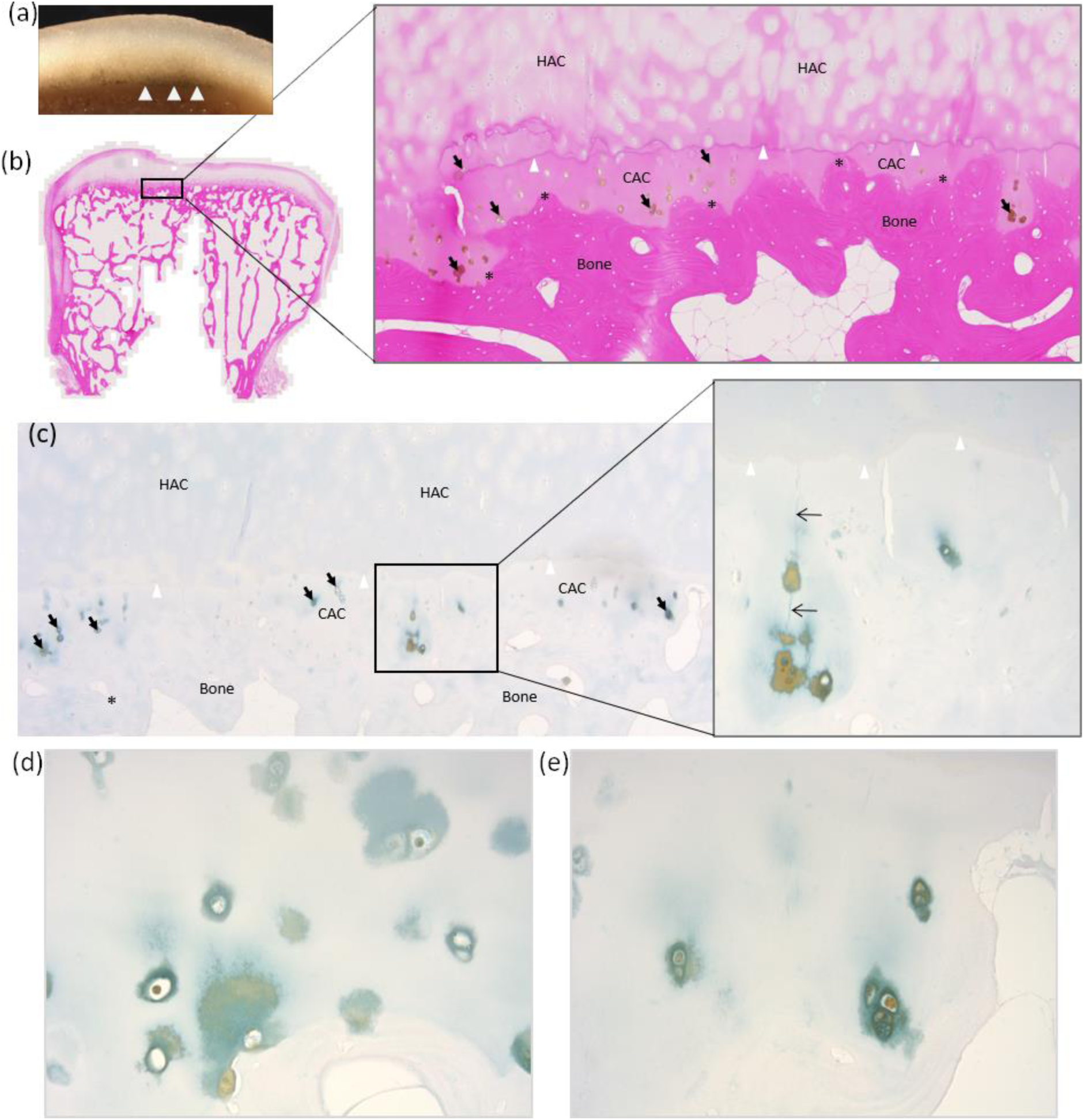
Ochronotic pigmentation of calcified articular cartilage. A cross section through the articular surface of the left radial head (post-fixation) is shown in (a), with the white arrow heads indicating pigmentation of the deep calcified articular cartilage adjacent to the subchondral bone that appears unpigmented. H&E staining of the radial head in is shown in (b), and Schmorl’s staining in (c). Pigmented chondrons, indicated by arrows, in (b) and (c) are observed in the calcified articular cartilage (CAC), deep to the tidemark indicated by white arrow heads, and not in the hyaline articular cartilage (HAC). Schmorl’s staining of the articular cartilage is shown in (c). Asterix’s indicate the position of the cement line separating the CAC from the underlying subchondral bone. The inset in (c), along with (d) and (e), show higher magnification images of pigmented chondrons within the CAC. The arrows in the inset of (c) show a pigmented line connecting adjacent pigmented chondrons and the superficial tidemark.

Perichondrium is associated with the periphery of the articular surface where cartilage is adjacent to bone, and dark pigmentation of perichondrium was observed. For example, the arrows in Figure 8c demonstrate dark, superficial pigmentation of the perichondrium on the capitate, at the junction of the cartilage and bone shaft. The radial head was cut into two segments and dark pigment was clearly observed on the superficial surface of the cartilage (see arrows Supplementary Figure 5a) at the edge of the articular surface towards the bone shaft, and not within the deeper cartilage which appears white on its cut surface. The periosteum is visible more distally, which appears whiter. The radial head was examined histologically at the periphery of the articular surface (dashed box area Supplementary Figure 5a), where pigmentation was observed at the superficial surface of the cartilage, indicated by arrows, which became fainter towards the main articular surface further away from the bone shaft (Supplementary Figure 5b). The pigment is located within the inner chondrogenic portion of the perichondrium, with the outer fibrous perichondrium appearing unpigmented. At junction of the perichondrium with the periosteum (Supplementary Figure 5c), a diffuse, granular-like pigmentation was observed within the perichondrium and adjacent hyaline articular cartilage. More distally, pigmentation within the periosteum was observed (Supplementary Figure 5d).

### 4.8 Lower limb joints

The hip and knee joints of both limbs were artificial and therefore not examined, except for a femoral head that was surgically removed approximately 3 years prior to death. The talus of the ankle joint was the most pigmented articular joint surface observed within the lower limb, with black to brown pigmentation across the talocrural joint surface, (Figure 8e). All other talar articular surfaces were pigmented, confined to the periphery of the articular surfaces. Pigmentation was observed on all other tarsal bone articular surfaces and the heads of the metatarsals, varying from light brown to dark brown/black, particularly at the edges (Figure 8f-g). An intensely pigmented inter-tarsal ligament insertion was observed on the lateral cuneiform which was black in colour (white arrow, Supplementary Figure 4a). The femoral head had intact cartilage that appeared to be pigmented across the centre of the articular surface (black arrows, Figure 8h) with very dark pigmentation of the perichondrium (white arrows) at the periphery of the articular surface adjacent to the bony neck of the femur. Pigmentation of chondrocytes within the CAC was observed histologically in the lateral cuneiform (not shown). Pigmentation was present on the periphery of the lateral cuneiform’s articular surface (see dashed box area of Supplementary Figure 4a) and was superficial. Histological examination of this region (Supplementary Figure 4b-c) showed more intense pigmentation at the periphery of the articular surface (left hand side of the image) that is located within the perichondrium.

### 4.9 Bone

Bone associated with the joints examined above grossly appeared normal. The periosteum was only investigated histologically in the radius at the junction with the articular cartilage perichondrium as already mentioned above. Trabecular bone of the radial head, capitate and lateral cuneiform examined histologically showed some pigmentation of osteocytes (Supplementary Figure 6c,g,f), observed only in a few regions on each of the sections examined, with the majority of osteocytes appearing unpigmented. In the ear ossicles (Figure 4), pigmented osteocytes were much more abundant than those observed in the limbs. Within the malleus (Figure 4a), pigmented osteocytes were more frequent in the handle and neck of the malleus. The pigmented osteocytes in the incus and stapes appeared to be distributed throughout the bone matrix (Figure 4b-c). Pigmentation of large, round cells assumed to be chondrocytes were observed in areas of trabecular bone matrix of the lateral cuneiform and radial head that did not stain strongly with eosin and did not have a lamellar structure (Supplementary Figure 6b,d), suggesting that these regions could be remnants of calcified cartilage or old woven bone that did not get replaced with lamellar bone. Schmorl’s staining confirmed pigmentation of the osteocytes and chondrocytes.

### 4.10 Other tissues and viscera

During life, pigmentation of the sclera in the temporal and nasal region was observed in both eyes and photographed, see Figure 10a,f. Post-mortem, histological sectioning of the right temporal sclera shows the presence of ochronotic pigment, with Schmorl’s staining (Figure 10a,c) showing a region of intense staining, that corresponds to an area of yellow coloured pigment in the H&E stained section (Figure 10d,e). Figure 10g shows a region of pigmented sclera stained with Schmorl’s stain, with the higher magnification image showing both diffuse and focal pigmentation, with the focal accumulations appearing more intense and yellow/brown in colour (Figure 10h).

**Figure 10.**
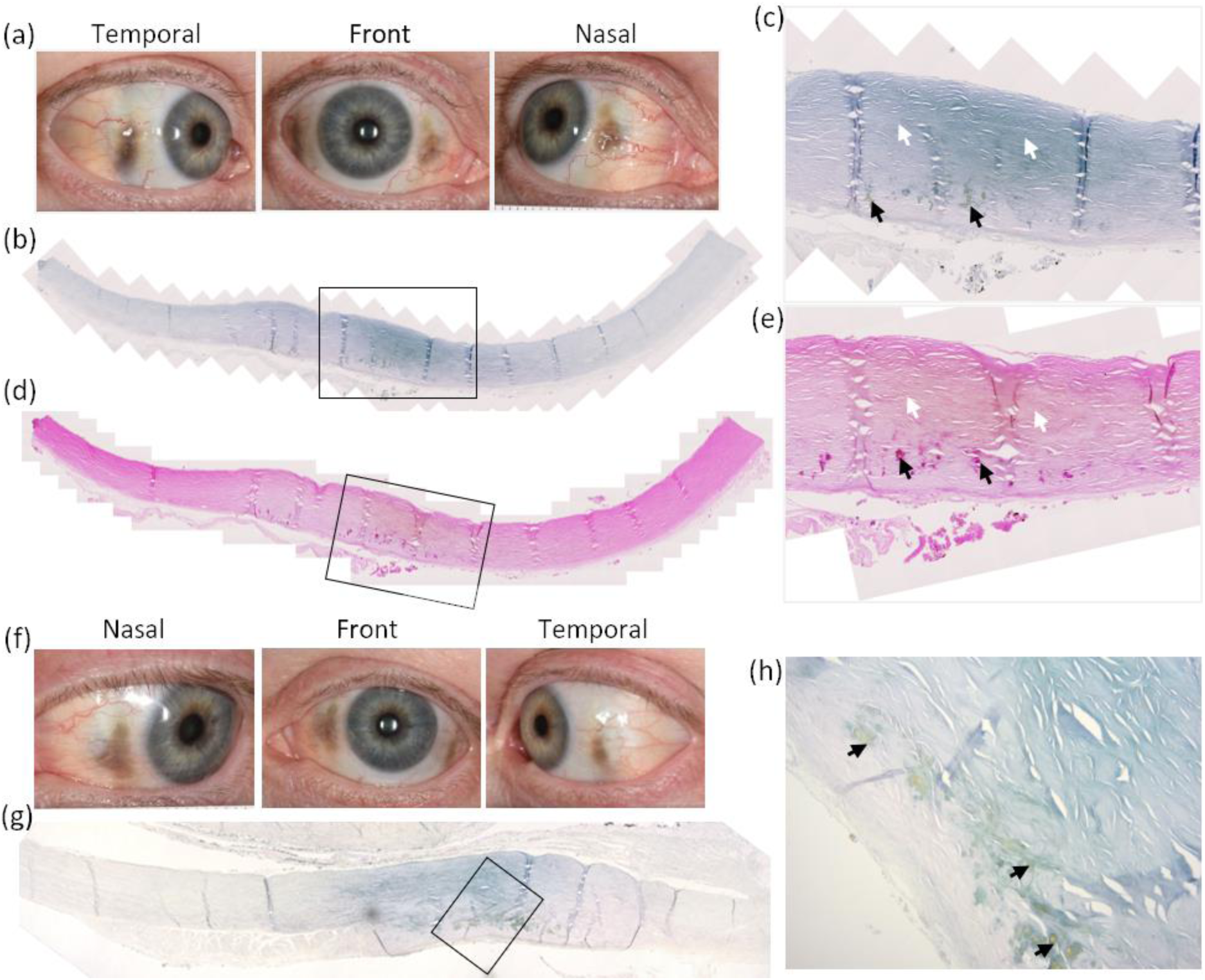
Ochronotic pigmentation of the sclera. Photographs of the right eye ante-mortem are shown in (a), showing ochronotic pigmentation in the nasal and temporal regions of the sclera. Histological sections of the right temporal sclera showing ochronotic pigmentation, stained with Schmorl’s stain in (b) and H&E in (d) are shown. The insets in (b) and (d) are shown in (c) and (e) respectively, with white arrows indicating diffusely pigmented areas, and black arrows indicating focal pigmentation. Photographs of the left eye ante-mortem are shown in (f), showing ochronotic pigmentation in the nasal and temporal regions of the sclera. A histological section of pigmented sclera from the left eye stained with Schmorl’s stain shown in (g), with a higher magnification image shown in (h) where diffuse pigmentation is observed throughout the sclera, and focal aggregates of pigment are indicated by arrows. Ante-mortem photographs of the eyes were taken 9 months prior to death.

Overall, the viscera appeared normal and healthy. Unless stated otherwise, the following viscera were examined grossly and histologically via H&E and Schmorl’s staining. The oesophagus, stomach, duodenum, jejunum, ileum, large intestine, liver, gallbladder, pancreas, spleen, kidney, bladder, ovary, skeletal muscle (gastrocnemius, adjacent to Achilles tendon), salivary glands (parotid, sublingual, submandibular), lacrimal gland and lymph nodes (carotid sheath and tracheal) appeared normal and unpigmented (Supplementary Figures 7 – 13). The stomach omenta and appendix appeared normal grossly but were not examined histologically. The lung parenchyma appeared unpigmented (Supplementary Figure 11a), with the cartilage of the bronchi and trachea described separately above. The uterus was only examined grossly and appeared unpigmented, with the only abnormal feature observed being a large fibrous cyst (fibroid) on the anterior part that upon transection appeared non-pigmented. The brain was removed from the skull and not examined further, but grossly appeared unpigmented.

## 5 Discussion

Here, an extensive anatomical and histological examination of tissues from an AKU individual, aged 60 years, has been carried out. Despite treatment for 7 years prior to death with nitisinone, the detrimental effect of pigmentation had already begun to take its toll on this individual; six major joint replacements had been carried out (both knees, hips and shoulders) and aortic valve function had declined requiring aortic valve replacement surgery, during which this individual passed away. It is evident that cartilage is very susceptible to pigmentation, particularly the calcified articular cartilage of joints and non-articular hyaline cartilage, where early pigmentation of the perichondrium is demonstrated. Furthermore, we show pigmentation of the ear ossicle synovial joints, the smallest in the body, the collagen-rich tympanic membrane and sclera, and show variable pigmentation of fibrocartilage at different anatomical sites.

The articular cartilage, particularly of large weight bearing joints, is the tissue most affected by ochronotic pigmentation, also confirmed in AKU mice (Preston et al., 2014, Hughes et al., 2021). Taylor et al. (2011) established the order of pigment deposition, beginning in the CAC associated with chondrons and territorial matrix, and then spreading to the HAC towards the superficial surface of the cartilage, encompassing the entire cartilage matrix. Here, we did not observe severe pigmentation of articular surfaces where the entirety of the articular surface is black, with most joints only showing mild pigmentation. The joints observed histologically were at an early stage of ochronosis, affecting only the CAC and not the HAC. Ochronosis was heterogenous across different joints, with the small joints observed here at an early ochronotic state. The most pigmented articular surfaces were the talus of the talocrural/ankle joint and the femoral head (removed 3.4 years antemortem for joint arthroplasty). Pigmentation of articular cartilage was greater in joints bearing more weight, for example the talus of the ankle joint compared to the distal humerus/proximal radius and ulna of the elbow, indicating that greater loading exacerbates pigmentation, with larger and more loaded joint replacements also having occurred in life. In the femoral head cartilage, centrally located pigmentation was observed, which decreased in intensity towards the periphery of the articular surface, as described by Taylor et al. (2011), then increased again with pigmentation of perichondrium at the junction of the articular surface with the bone (perichondrium is discussed below).

Here, we examined small articular joint surfaces with a mild ochronotic phenotype such as the distal humerus, carpal and tarsal bones. The pigmentation was focussed at the periphery of articular surfaces, rather than located more centrally. The superior articular surface of the talus (talar dome), at the talocrural/ankle joint did show a wider area of pigmentation than other smaller bones which is not surprising as the talar dome carries approximately 77-90% of the load carried across the tibial-talar joint (Brockett and Chapman, 2016), but was more intense more anteriorly and posteriorly, rather than centrally. The talar dome pigmentation therefore does not appear to be in the area of greatest load or contact area, which is described as centrally and laterally by Wan et al. (2006). It may be that centrally located pigment, as described by Taylor et al. (2011) in hips and knees, is a feature of joints with more advanced ochronosis. Here we only observed minimal pigment confined to the CAC layer of cartilage, which may not have been visible at the centre of the articular surfaces where articular cartilage is thicker and masked by overlying unpigmented HAC, and therefore only visible at the periphery where it is thinner (Fox et al., 2008, Zumstein et al., 2014, Cohen et al., 1999). A limitation of our examination may be that most articular surfaces were not viewed in cross-section; the radial head did demonstrate visible pigmentation in the CAC in cross-section, that appeared greater in the middle compared to the edges more laterally. Furthermore, it has been previously shown that AKU individuals do not have typical gait (Barton et al., 2015, Shepherd et al., 2022). Due to having arthropathy and arthroplasty in other joints, such as both knees and both hips here, and having ochronosis in other musculoskeletal tissues affecting function, cartilage is likely loaded abnormally which may affect the location of pigment.

The tidemark, or current mineralising front as it is also known, is the interface between the HAC and CAC. Tidemark duplication was observed histologically in several articular cartilage samples, with pigmented chondrons located between the two tidemarks, with three observed in some cases. Tidemark duplication or advancement is often observed in osteoarthritis (Oettmeier et al., 1989), with a dual tidemark associated with a higher OARSI grade in human joints (Palmer et al., 2014). Tidemark multiplication however can also be considered a normal feature of articular cartilage (Boyde, 2021, Oegema et al., 1997, Lane and Bullough, 1980). Palmer et al. (2014) suggest that tidemark advancement occurs at an early stage of OA and may perpetuate further osteoarthritic changes in the HAC in a “bottom-up” mechanism. We know that AKU joints become osteoarthritic due to pigment deposition in the CAC before the HAC also in a “bottom-up” manner, therefore it is plausible that the tidemark multiplication observed here is pathological. Interestingly, pigmented chondrons were observed between two tidemarks within CAC; whether these chondrons become pigmented after tidemark advancement/matrix calcification or before when they were still HAC chondrocytes is unknown. We speculate that it is likely after tidemark advancement, as no pigmented chondrons were observed in the adjacent HAC, suggesting that the environment of chondrons within CAC promotes pigmentation.

It is possible that the low oxygen tension (Brighton et al., 1971, Zhou et al., 2004) and acidic pH, reported to be as low as 6.6 – 7.2 (High et al., 2019, Urban et al., 1993, Gray et al., 1988, Razaq et al., 2003), of cartilage may promote pigmentation, however other factors must be involved to explain why CAC pigments before the neighbouring HAC. CAC is very understudied, as reviewed by Evans and Pitsillides (2022). The importance of CAC towards joint pathologies such as OA has begun to be uncovered, with examination of osteoarthritic mice also showing early pathological changes in CAC chondrocytes that precede OA (Herbst et al., 2023, Madi et al., 2020, Staines et al., 2016). Interestingly, it was observed that within the CAC, the most intensely pigmented chondrons were situated in the deepest layers of the CAC where the undulating cement line is deeper, and therefore a further distance from the tidemark. Whether this is significant towards the pathological process of ochronosis is unknown but may warrant further investigation with examination of samples from different AKU donors.

In addition to CAC pigmentation, we present other skeletal connective tissues and cells that pigment, such as perichondrium, non-articular hyaline cartilage, fibrocartilage and bone osteocytes. The hyaline costal cartilage was extremely pigmented, grossly appearing black in colour, along with the respiratory cartilage of the larynx and upper trachea. Histologically, pigmentation with non-articular hyaline cartilage appeared diffuse rather than focal, such as within the tracheal ring and costal cartilage. None of the joints examined histologically here exhibited HAC pigmentation, therefore we were unable to compare articular HAC pigment distribution to the costal and tracheal cartilage. Taylor et al. (2011) however described HAC pigmentation as blanket rather than focal, with more widespread pigmentation observed than in CAC. We suggest that the pigment HAC of joints and non-articular hyaline cartilage is similar and diffusely spread, whereas pigmentation of CAC appears much more focal and related to the chondrocytes. Taylor et al. (2011) report that once pigmentation begins in HAC, it progresses and spreads rapidly throughout the matrix, faster than the underlying CAC. The black appearance of the non-articular hyaline cartilage in this examination (i.e. trachea, costal) would support this, compared to the less intense pigment observed in joints. Why the non-calcified hyaline cartilage of joints is initially resistant to pigmentation is not known, and we suggest that changes to the matrix may occur that then allows this rapid pigmentation.

Perichondrium is the lining surrounding non-articular hyaline and elastic cartilage. Although articular cartilage is devoid of perichondrium on the main articular surface, it is located at the periphery of articular cartilage, adjacent to the bone, where it blends with periosteum. Perichondrium is composed of an outer fibrous layer with fibroblasts, with an inner chondrogenic or cellular layer that can give rise to new cartilage cells via chondroblasts during appositional growth of cartilage (Ross and Pawlina, 2015). Perichondrium pigmentation has been described before, but with no discussion around its presence. Pigmentation of the perichondrium of respiratory hyaline cartilages (tracheal, bronchial, thyroid, cricoid) and costal cartilage is described, with a few available images showing it to be associated with the inner perichondrium layer as we have described, spreading into the hyaline cartilage matrix (Helliwell et al., 2008, Lichtenstein and Kaplan, 1953, Gaines, 1989, Nishimori et al., 1970). We showed in multiple cartilages that perichondrium pigmentation was more prominent than that of the hyaline cartilage matrix that it surrounds, comparable to a description of respiratory cartilage by Gaines (1989). The external auditory meatus, nasal septum and secondary bronchial cartilage however only showed perichondrium pigmentation, suggesting that pigment is deposited in the perichondrium before the cartilage matrix. Examination of the costal cartilage did however show more intense pigmentation in the hyaline cartilage compared to the perichondrium, therefore we would hypothesize that although perichondrium pigmentation precedes cartilage pigmentation in these tissues, it proceeds more rapidly in the hyaline cartilage, similar to the HAC of joints as discussed above. Pigmentation of perichondrium related to joints has been reported by Lichtenstein and Kaplan (1953), although they did not state the joints examined.

Intense pigmentation was observed associated with chondrocytes within joints. Additionally, pigmentation was observed that appears to connect two chondrocytes (inset of Figure 9c), with the pigmented “line” also extending from the more superficial chondrocyte to the tidemark. This could potentially be evidence of chondrocyte connectivity, which has been a debatable topic over the years. Chondrocyte connectivity has been observed in bovine cartilage using an optical clearance method to visualise cytoskeletal components in load bearing femoral articular cartilage (Neu et al., 2015). A group investigating cytoplasmic processes in human cartilage showed that fine processes were observed extending away from 40% of chondrocytes in human tibial cartilage at all stages of cartilage degeneration, with variable presence, numbers per cell (1-9) and length (1-40µm) (Bush and Hall, 2003, Murray et al., 2010). They also showed that there was reduced pericellular collagen type VI coverage in cells with long processes and proposed that early pericellular matrix weakening could lead to loss of chondrocyte-matrix interactions and development of micro-fissures through which a cytoplasmic process could develop. It is possible that ochronotic pigment within chondrons in AKU cartilage also leads to weakening of the pericellular matrix and abnormal cell-ECM interactions. Cytoplasmic processes up to 30µm in length have been identified in 20-45% of OA femoral cartilage compared to <10% in non-OA, correlating with the extent of histological OA degradation (Holloway et al., 2004). It is possible that cytoplasmic processes could be a pathological feature caused by pigmentation in AKU in which OA manifests, however Holloway et al. (2004) report that they did not observe any cytoplasmic processes in histological sections (5µm thick). Cell projections however have been identified in human and boar articular cartilage sectioned at 4µm with standard optical microscopy, confirmed via confocal and electron microscopy to be long cellular projections from 5 to 150µm in length (Mayan et al., 2015). Whether we have observed examples of cytoplasmic processes here cannot be conclusively stated; immunohistochemical staining of cytoplasmic proteins or confocal laser scanning microscopy in the future would be required.

It was considered that the chondrocyte extensions observed here were pigmented primary cilia. Primary cilia are found on chondrocytes of articular cartilage, essential for mechanotransduction, with an altered response to mechanical stimuli in their absence, and differences found in healthy and osteoarthritic cartilage (Ruhlen and Marberry, 2014, Wann et al., 2012, McGlashan et al., 2010). Although we showed that the extension was projecting in the direction of the articular surface/subchondral bone as described by Farnum and Wilsman (2011), primary cilia only 1-2µM in length (Meng et al., 2023), much smaller than the structure we observed. Primary cilia have been investigated in an AKU cell culture model (Gambassi et al., 2017), in which they were reduced in length compared to non-AKU conditions, with an alteration of intracellular signalling. With loss of primary cilia leading to OA in a mouse model (Irianto et al., 2014), and AKU being a rare form of OA, the effect of HGA/pigmentation on the cell, the ECM and cell-ECM interactions is likely to have a negative effect on cartilage homeostasis and is worthy of future investigation.

We demonstrate here pigmentation of CAC within the synovial joints between the ear ossicles and also of the osteocytes (bone is discussed below). Along with pigmentation of articular cartilage, pigmentation was also observed in the tympanic membrane and perichondrium associated with the tympano-mallear connection. Pigmentation was observed in the dense connective tissue of the tympanic membrane which blends with the lamina propria layer, formed mostly of type I collagen. The tympanic membrane and synovial joints of the ear ossicles would be under constant strain from sound vibrations, therefore although they are small, it’s not surprising to find pigmentation here. Previous biomechanical research has shown that AKU articular cartilage is stiffer than non-AKU and OA cartilage (Taylor et al., 2011). It may be that all pigmented tissue become stiffer upon pigmentation, which may have functional consequences. Stiffening of the tympanic membrane could therefore be occurring in AKU patients, which could have an impact on hearing.

Tympanic membrane discolouration is well documented in the literature (Sanji et al., 2011, Sagit et al., 2011, Ozer Ozturk et al., 2022, Pau, 1984, Steven et al., 2015, Al-Shagahin et al., 2019). The ear ossicles have never been described to our knowledge. We would propose that pigmentation of the tympanic membrane and synovial ear ossicle joints would cause conductive hearing loss affecting sound conduction through the outer and middle ear cavities, with abnormal tympanograms. Mild to moderate hearing loss in AKU has been shown in the literature, with an incidence of 35% (n=20) and 44% (n=16) in two studies, at an average age of 63 and 54 years respectively (Steven et al., 2015, Al-Shagahin et al., 2019). Surprisingly, the majority of the hearing loss was sensorineural in these studies, with just a few cases reporting mixed hearing loss (combination of conductive and sensorineural) (Sagit et al., 2011, Ozer Ozturk et al., 2022, Pau, 1984), all with normal tympanograms, suggesting that the inner ear/central nervous system may in fact be involved. Cochlear function could be affected by HGA and/or pigmentation, however this has never been assessed. The absence of acoustic reflexes in the two cases reporting mixed frequency hearing loss with normal tympanograms (Sagit et al., 2011, Ozer Ozturk et al., 2022) however could point to stiffness/abnormal function of the ear ossicle chain that prevents the acoustic reflex acting on the tympanic membrane, or perhaps even stiffness of the tensor tympani/stapedius tendons themselves. The mechanism in which hearing is impaired in AKU remains to be established.

Whilst most connective tissues, particularly those that are collagen rich, are susceptible to pigment in AKU, the collagen I-rich bone matrix appears to be protected. Grossly, we did not observe pigmentation of bone, and only observed it in periosteum, agreeing with previous macroscopic observations (Lichtenstein and Kaplan, 1953, Helliwell et al., 2008). Histologically, we observed pigmentation of osteocytes observed in trabecular bone of appendicular joints and the ear ossicles via histology, but not within the bone matrix, also observed by Taylor et al. (2011). Osteocyte pigmentation in the trabecular bone from limbs was scarce in comparison to the numerous pigmented osteocytes observed throughout the bone matrix of the ear ossicles. The unique function and composition of the ear ossicles may explain why they are more susceptible to pigmentation.

The ear ossicles reach their final shape and size by birth which is unusual for most bones and are composed exclusively of compact bone, with no trabecular bone, and are formed of a mixture of lamellar and woven bone (Hassmann and Chodynicki, 1978, Duboeuf et al., 2015). Two human ear ossicle studies reported low levels of bone remodelling, with almost none observed (Duboeuf et al., 2015, Chen et al., 2008). Two other studies showed that empty osteocyte lacunae increase dramatically from birth via apoptosis in human ear ossicles, in addition to the number of degenerating osteocytes, with no apoptosis observed in older age (i.e. 60 years), with the hypothesis that osteocyte apoptosis is a programmed phenomenon whereby cell death inhibits bone remodelling, ensuring that the ear ossicles acquire the structural stability they need for sound transmission (Marotti et al., 1998, Rolvien et al., 2018). Here, the lack of bone turnover would result in older osteocytes that have had longer exposure to HGA and more opportunity to become pigmented, compared to osteocytes of newly formed bone. Additionally, it has been shown that osteocyte lacunae of ear ossicles are hyper-mineralised/micropetrotic, with most of these changes occurring within the first year of life (Rolvien et al., 2018), occurring after osteocyte cell death (Milovanovic and Busse, 2023). This suggests that many, if not most of the pigmented osteocytes observed here within the ear ossicles may be dead. Whether they become pigmented before or after death cannot be deduced via our observations, as we did not look at viability, the tissue was decalcified for examination.

At a few sites within the trabecular bone, pigmented osteocytes were observed that were surrounded by fully mineralised, lamellar bone. These osteocytes tended to be clustered together, suggesting an area of older bone with longer lived osteocytes. Additionally, there were areas within the trabecular bone network where pigmentation appeared to be associated with rounder, chondrocytic-looking cells surrounded by a non-mineralised matrix that stained weakly with eosin and lacked a lamellar structure. This matrix appears to be remnant calcified cartilage that has not been replaced by lamellar bone, and is susceptible to pigmentation.

So far, hyaline cartilage and elastic cartilage pigmentation has been demonstrated. The third major type of cartilage, fibrocartilage, is susceptible to pigmentation, although high quality histology images are not often reported. Fibrocartilage is found at many sites across the body, lacks a perichondrium and functions to resist compression, shear force and tensile resistance. Fibrocartilage is composed of dense fibrous tissue with chondrocytes and fibroblasts, that are surrounded by a matrix that contains type I collagen in addition to type II collagen. Previously, pigmentation has been described in intervertebral discs (Helliwell et al., 2008, Galdston et al., 1952), the pubic symphysis (Helliwell et al., 2008, Lichtenstein and Kaplan, 1953) and knee joint menisci (Nag et al., 2013, Xu et al., 2015, Rajani et al., 2021). Mouse studies have shown that chondrons of calcified fibrocartilage pigment at tendon and ligament entheses, in addition to knee joint menisci chondrons (Hughes et al., 2021), however pigmentation was not found within the fibrocartilaginous intervertebral discs of mice.

Pigmentation of the pubic symphysis disc was observed here, and appeared patchy, with focal deposits associated with cells, agreeing with previous reports of human pubic symphysis pigmentation. To our knowledge, we are the first to examine the TMJ in AKU and were surprised by the lack of pigmentation. The TMJ in previous investigations may not have been described because it is unaffected by AKU, or because it was not examined. Although the TMJ articular surfaces of the mandible and temporal bone macroscopically appeared unpigmented, we cannot say with certainty that pigmentation was not present as it was not examined histologically. Future work examining the TMJ articular surfaces would be interesting due to the articular cartilage composition being both fibrocartilaginous and hyaline-like, with type I collagen and type II collagen both present (Mizoguchi et al., 1996, Wang et al., 2009, Delatte et al., 2004, Teramoto et al., 2003). Although its biomechanics are complex, the TMJ is considered a loaded joint. With the TMJ fibrocartilage articular disc having an extracellular matrix composition of type I collagen and elastin, with only trace amounts of type II collagen and other collagens (Runci Anastasi et al., 2021, Detamore and Athanasiou, 2003, Landesberg et al., 1996), suggesting it is subjected to high tensile strain and deformation, it was very unexpected to find no evidence of pigmentation. We have only been able to study the TMJ in one AKU individual, therefore more evidence in other individuals would be desirable to confirm if this is a commonly unaffected tissue or not. However, due to this tissue being extremely inaccessible and unlikely to be subjected to joint arthroplasty, it is unlikely another human TMJ from an AKU individual will be available to study soon. AKU mice therefore present as the best opportunity to understand the pathology (or lack of) within the TMJ in AKU, in addition to studying other fibrocartilage such as the sternoclavicular and pubic symphysis articular discs which have not previously been examined in mice (Hughes et al., 2021).

Scleral pigmentation is a well-recognised sign of AKU. The sclera is a dense fibrous tissue predominantly formed from type I collagen and small amounts of type III collagen (Keeley et al., 1984), embedded in a proteoglycan matrix. We show the first histology of pigmented sclera showing both diffuse and focal aggregates of pigmentation. Grossly, the pigmentation appears to be deep to the blood vessels situated within the episcleral layer, and therefore is in the thickest, deeper stromal layer, confirmed by histology. The nasal and temporal scleral pigmentation observed here is well reported in AKU and has been assessed routinely in some cohorts of AKU patients (Ranganath et al., 2020b). A systematic review of 36 case reports found the most frequent AKU sign was symmetric pigmentation in the sclera in 82% of patients, located within the palpebral fissure (Lindner and Bertelmann, 2014). Where limbal pigmentation was noted, it was located at the level of Bowman’s membrane or adjacent to it, with an “oil drop” like appearance. Conjunctival pigmentation was observed in 60% of the cases. Interestingly, the majority of pigment was found to reach from the insertion of the rectus muscles towards the pars plana (located between the ciliary body and retina, near the junction of the sclera and iris). It has previously been suggested that both UV light damage and stress arising from muscle contraction may be damaging factors that lead to scleral/conjunctival pigmentation (Ranganath et al., 2019). Although UV light could be a contributing factor, it wouldn’t explain the localisation of pigment to the nasal and temporal sclera only. Stress from nearby extraocular muscle contraction may explain the occurrence of pigmentation, perhaps from the medial and lateral recti muscles, however it does not explain lack of pigmentation superiorly and inferiorly, where the superior and inferior recti muscles, and the oblique muscles attach. It is possible that the deeper ciliary body muscles could be causing tension on the sclera. Why pigmentation consistently affects the temporal and nasal sclera is both intriguing and currently unknown.

As expected, pigmentation was observed in collagen-rich connective tissues with most soft and visceral tissue appearing normal and unpigmented. The distribution of pigmentation was very heterogenous both across and within the connective tissues, with vast differences in pigment intensity. This heterogeneity however is not a new phenomenon; even between siblings carrying the same genetic HGD variants, considerable variability in the AKU disease phenotype has been observed (Zatkova et al., 2022, Vilboux et al., 2009). Heterogeneity of ear cartilage pigmentation is reported, and has been described as localised and diffuse, and not uniform across the tissue (Al-Shagahin et al., 2019, Ranganath et al., 2020a). Zatkova et al. (2022) suggest other factors such as genetic, biomechanical and environmental factors modify the disease, causing accelerated pigmentation and tissue damage in some patients and not others.

The ochronosis observed in this dissection is much less than that observed by Helliwell et al. (2008), however the individual was 74 years old and had also not recieved HGA-lowering treatment. The individual here was 60 years old and had received nitisinone for 7 years prior to death. There could have been some reversal of ochronosis on nitisinone therapy, as previously described by Ranganath et al. (2020a), who showed a decrease in ochronosis scores of externally visible eye and ear pigment and histological assessment of ear biopsies. This would suggest that pigmentation may have been reversed in scleral and elastic auricular cartilage. The earlier work in AKU mice however demonstrated that cartilage pigmentation could only be halted with nitisinone treatment, and not reversed (Keenan et al., 2015). More studies are needed to investigate the potential mechanism of pigment removal, and must consider factors such as tissue type, location and function; it is possible that scleral/elastic cartilage pathology differs to hyaline joint cartilage and are therefore not comparable. Care must also be taken in interpretation due to the variation in disease severity between individuals.

The heterogeneity of pigmentation may provide an insight into its pathophysiology. It is clear that pigmentation favours connective tissues, particularly those that have a high fibrillary collagen content, namely collagen types I and II. In addition to different functions and tissues compositions, tissues also age and degenerate at different rates. The exposed collagen hypothesis of ochronotic pigmentation suggests that pigmentation requires changes in the extracellular matrix to occur (Gallagher et al., 2016), with pigmentation taking weeks to become visible histologically in mice (Keenan et al., 2015, Hughes et al., 2021) and years to become visible externally in humans. Human AKU joints after arthroplasty also exhibit areas not affected by pigmentation (Taylor et al., 2011). Whilst there is some evidence that points towards collagen as the binding site of HGA/pigment (Chow et al., 2011, Taylor et al., 2011), it has not been definitively proven. The exposed collagen hypothesis suggests that loss of protective molecules that decorate collagen such as proteoglycan and glycosaminoglycans allows pigmentation to occur, with changes in extracellular matrix occurring naturally through normal ageing and degenerative processes, overloading and trauma. It is also likely that pigmentation of cells and their surrounding matrix then influences metabolism and activity, however these affects are not known.

There are many fundamental questions that remain unanswered in AKU. The chemical structure of ochronotic pigment must still be elucidated as reviewed by Ranganath et al. (2019), along with the binding site within tissue extracellular matrix. There is therefore merit in identifying the connective tissue niche and microenvironment that pigment favours, as its binding sites to protein may reveal information about its structure and reactivity. Whilst the current manuscript has not investigated pigmentation at the nanoscale level, histological analysis is a step forward. Here we confirm that within osteochondral tissues, calcified articular cartilage is very susceptible to pigmentation, and show that perichondrium pigments early in both articular and non-articular cartilage, and before non-calcified hyaline cartilage. We also show the heterogeneity of pigmentation both within and between different tissues and suggest that studying the fibrocartilaginous disc of the TMJ may provide insights into factors that are protective against ochronotic pigmentation. Whether or not pigmentation is able to reverse within joint tissue when treated with nitisinone requires further investigation.

## 6 Acknowledgements

We would like to acknowledge the generous gift of body donation from the donor and their family, allowing us to study this rare disease.

## 7 Conflict of interest

The authors have no conflicts of interest to declare.

## 8 Author contributions

JHH, APB, NPT, CMT, PDR, GC and AP were involved in tissue dissection and analysis. JHH and APB wrote the manuscript. All authors contributed to critical revision and approval of the manuscript. JAG and LRR conceived the study design.

## Notes

### Competing Interest Statement

The authors have declared no competing interest.

