## Supplementary for "An anatomical investigation of alkaptonuria: Novel insights into ochronosis of cartilage and bone"

### 1 Supplementary information

#### 1.1 Tissue processing

Three different protocols were used for tissue processing. Visceral/soft tissues were processed using protocol 1, and decalcified bone and cartilage were processed using protocol 2, see Supplementary Table 1. Eye tissue was processed using a different protocol, see Supplementary Table 2. Paraffin wax was at 62°C.

Supplementary Table 1. Tissue processing steps for soft and hard tissues.

| Step | Chemical | Protocol 1 | Protocol 2 |
|---|---|---|---|
| 1 | 70% ethanol | 30 mins | 15 mins |
| 2 | 90% ethanol | 30 mins | 1 hour |
| 3 | 100% ethanol | 10 mins | 1 hour |
| 4 | 100% ethanol | 10 mins | 1 hour |
| 5 | 100% ethanol | 10 mins | 1 hour |
| 6 | 100% ethanol | 10 mins | 1 hour |
| 7 | Xylene | 30 mins | 1 hour |
| 8 | Xylene | 30 mins | 1 hour |
| 9 | Paraffin wax | 2 hours | 2 hours |
| 10 | Paraffin wax | 2 hours | 2 hours |

Supplementary Table 2. Tissue processing steps for eye tissue.

| Step | Chemical | Protocol 3 |
|---|---|---|
| 1 | 70% ethanol | 40 mins |
| 2 | 90% ethanol | 40 mins |
| 3 | 100% ethanol | 40 mins |
| 4 | 100% ethanol | 40 mins |
| 5 | 100% ethanol | 1 hour 30 mins |
| 6 | 100% ethanol | 2 hours |
| 7 | Xylene | 1 hour |
| 8 | Xylene | 1 hour |
| 9 | Xylene | 2 hours |
| 10 | Paraffin wax | 1 hour 15 mins |
| 11 | Paraffin wax | 1 hour |
| 12 | Paraffin wax | 3 hours |

#### 1.2 Staining

Schmorl's staining was carried out using a freshly made incubating solution, that contained 1% ferric chloride (freshly made) and 1% potassium ferricyanide (stored at 4°C for up to 1 month) in distilled water. The nuclear fast red counterstain contained 0.1% nuclear fast red and 5% aluminium sulfate in distilled water. Slides were cleared in xylene twice (4 minutes each) and rehydrated using an ethanol series (100%, 90% and 70%, each for 2 minutes) and placed in water for 1 minute. Bone and cartilage slides were then placed in the incubating solution for approximately 10 minutes, with the viscera incubating for approximately 7.5 minutes. All slides were then washed in running tap water for 3 minutes, followed by immersion in 1% acetic acid for 5 minutes. Slides were counterstained for approximately 2 mins in nuclear fast red, and washed in running tap water, with agitation, for 5 minutes. All slides were then dehydrated in an ethanol series (70%, 90% and 100%, each for 2 minutes), and cleared in two changes of xylene for 3 minutes each. All slides were cover slipped using DPX.

H&E staining was carried out by clearing the slides in xylene twice (2 minutes each) and rehydrated using an ethanol series (100%, 90% and 70%, each for 1 minute) and placed in running tap water for 1 minute. Slides were then immersed in Harris Haematoxylin (filtered before use) for 5 minutes, washed in running tap water for 1 minute, and dipped in 1% acid alcohol for 15 seconds. Slides were then washed in running tap water for 5 mins, followed my immersion in alcoholic Eosin Y for 3 minutes and returned to running tap water for 15 seconds. Slides were then dehydrated in an ethanol series (70%, 90% and 100%, each for 30 seconds), and cleared in two changes of xylene for 2 minutes each. All slides were cover slipped using DPX.

#### 1.3 Histology of bone and cartilage

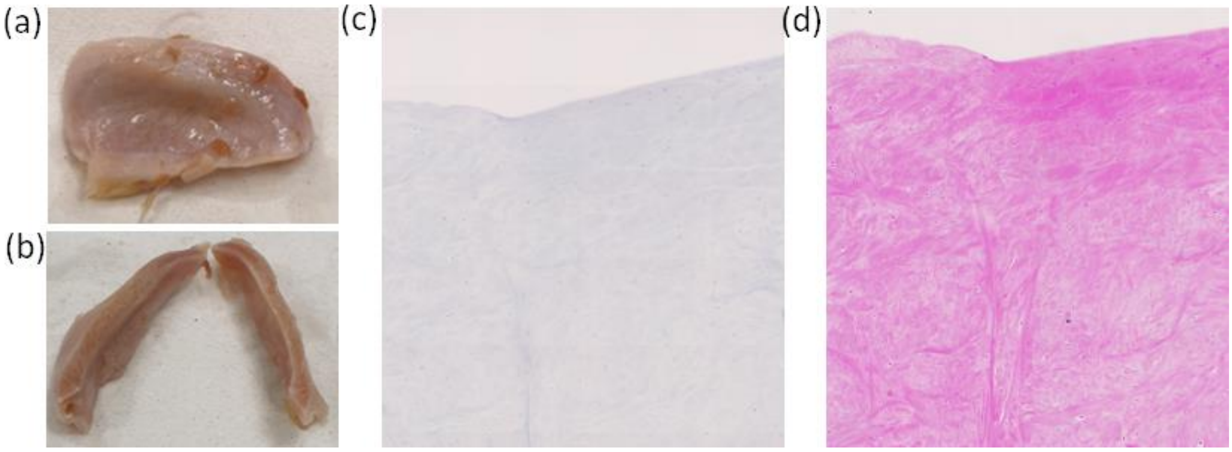


**Supplementary figure 1. Normal appearance of the temporomandibular joint articular disc.** The articular disc of the temporomandibular (TMJ) disc is shown in (a) and is bisected in (b), with no ochronotic pigmentation observed. Schmorl's and H&E staining of the TMJ articular disc is shown in (c) and (d) respectively, again with no pigmentation identified.

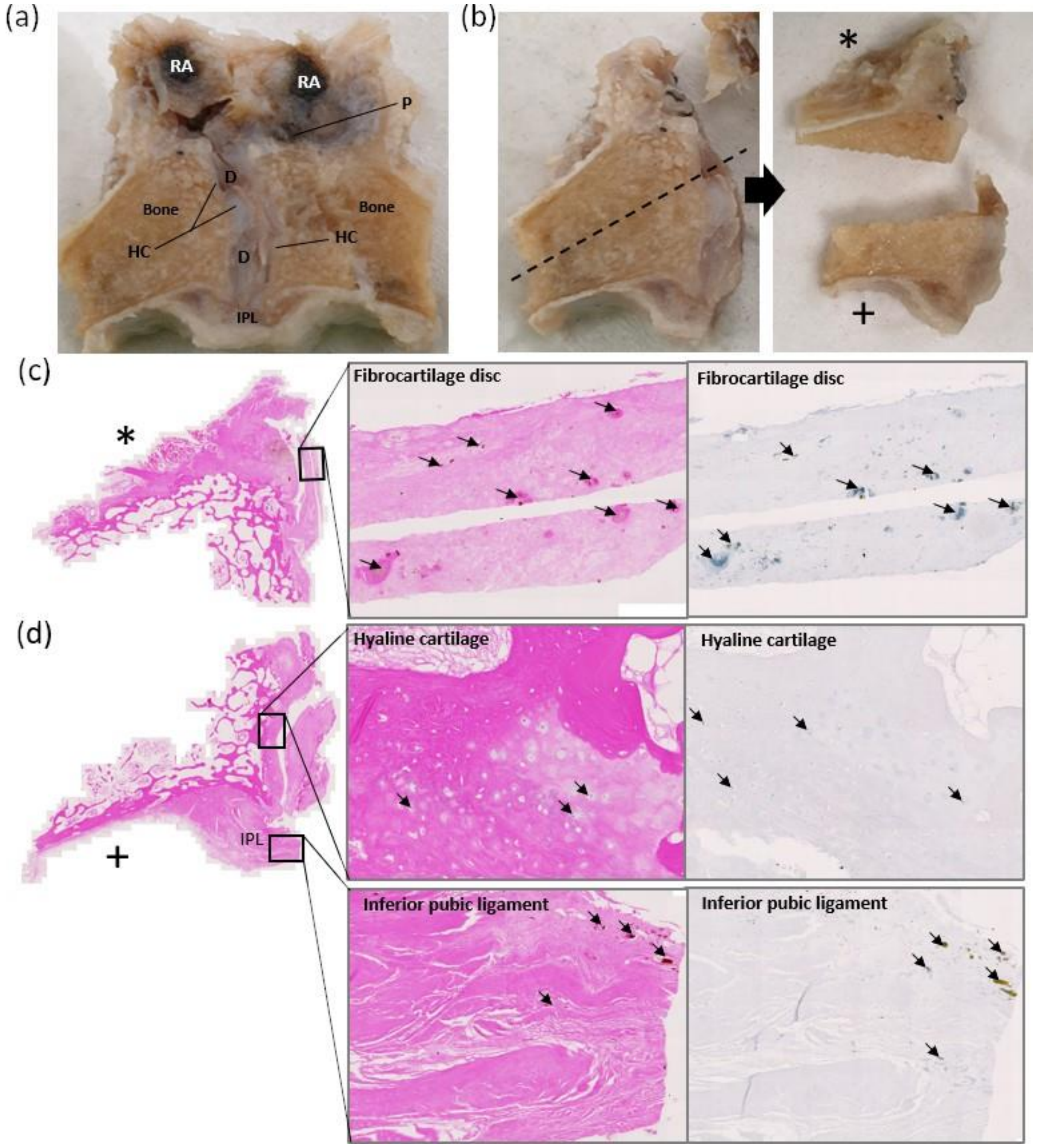


**Supplementary figure 2. Gross and histological ochronotic pigmentation of the pubic symphysis.** A coronal cross-section through the pubic symphysis joint (post-fixation) is shown in (a). The trabecular bone can be observed, with hyaline cartilage (HC) lining the joint surface, with a fibrocartilaginous inter-pubic disc (D) observed in the middle of the joint, with no ochronotic pigmentation observed macroscopically. Inferiorly, the inferior pubic ligament (IPL) can be observed, which appears grossly normal. The heavily pigmented tendons of rectus abdominis (RA) can be observed superior to the pubic symphysis joint, with a small of part of pyramidalis (P) tendon that is also pigmented. The segment of the pubic symphysis that was examined histologically is shown in (b), and cut into two separate parts; the top part (indicated by *) is shown in row (c) and the bottom part (indicated by +) is shown in row (d). H&E staining in (c) shows the fibrocartilaginous disc has a cleft, with higher magnification image showing areas of pigmentation indicated by arrows, that was also observed with Schmorl's staining (far right of row (c)). Row (c) shows H&E staining, with the insets showing the hyaline cartilage and inferior pubic ligament, with Schmorl's staining of the same areas also shown. Within the hyaline cartilage, minimal pigmentation was observed within a few chondrocytes, and was not very dark or intense with Schmorl's staining. The inferior pubic ligament showed areas of dark pigmentation, indicated by arrows.

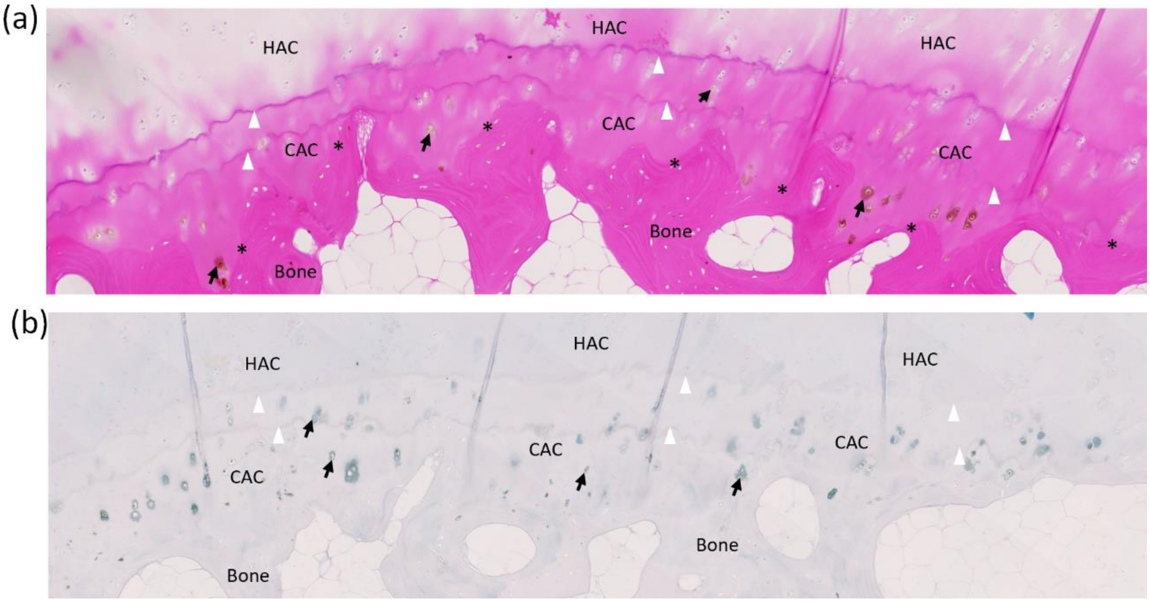


**Supplementary figure 3. Histological pigmentation of the capitate articular cartilage.** Histology of the left capitate articular cartilage is shown in (a) and (b), stained with H&E and Schmorl's stain respectively. In both (a) and (b), no pigmentation of the hyaline articular cartilage (HAC) is observed, with all pigmentation confined to the calcified articular cartilage (CAC), deep to the tidemark (denoted by white arrow heads). Example of pigmentation associated with chondrons are shown by black arrows. The cement line, the boundary between the CAC and bone, is indicated by asterix's in the H&E in (a).

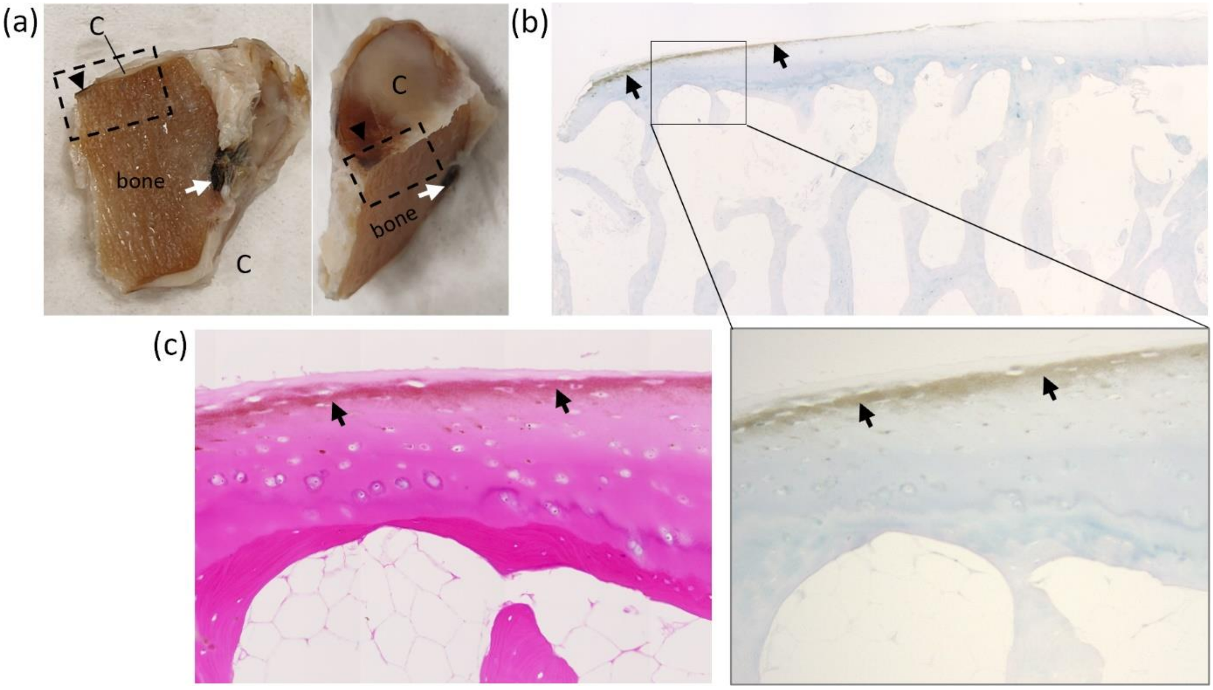


**Supplementary Figure 4. Pigmentation of lateral cuneiform articular cartilage, inter-tarsal ligament and the perichondrium**. Part of the lateral cuneiform bone (post-fixation) is shown in (a) with the white arrow pointing to an inter-tarsal ligament insertion that was intensely pigmented, and the black arrows heads indicating pigmentation of the surface of the articular cartilage. The dashed region of the articular cartilage in (a), shown from two viewpoints, is shown histologically in (b), stained with Schmorl's stain. The pigmentation that could be observed grossly associated with the articular cartilage in (a), is shown to be pigmentation of perichondrium in (b), indicated by arrows. H&E staining of the area shown in the inset in (b), is stained with H&E, where perichondrium pigmentation was observed (c).

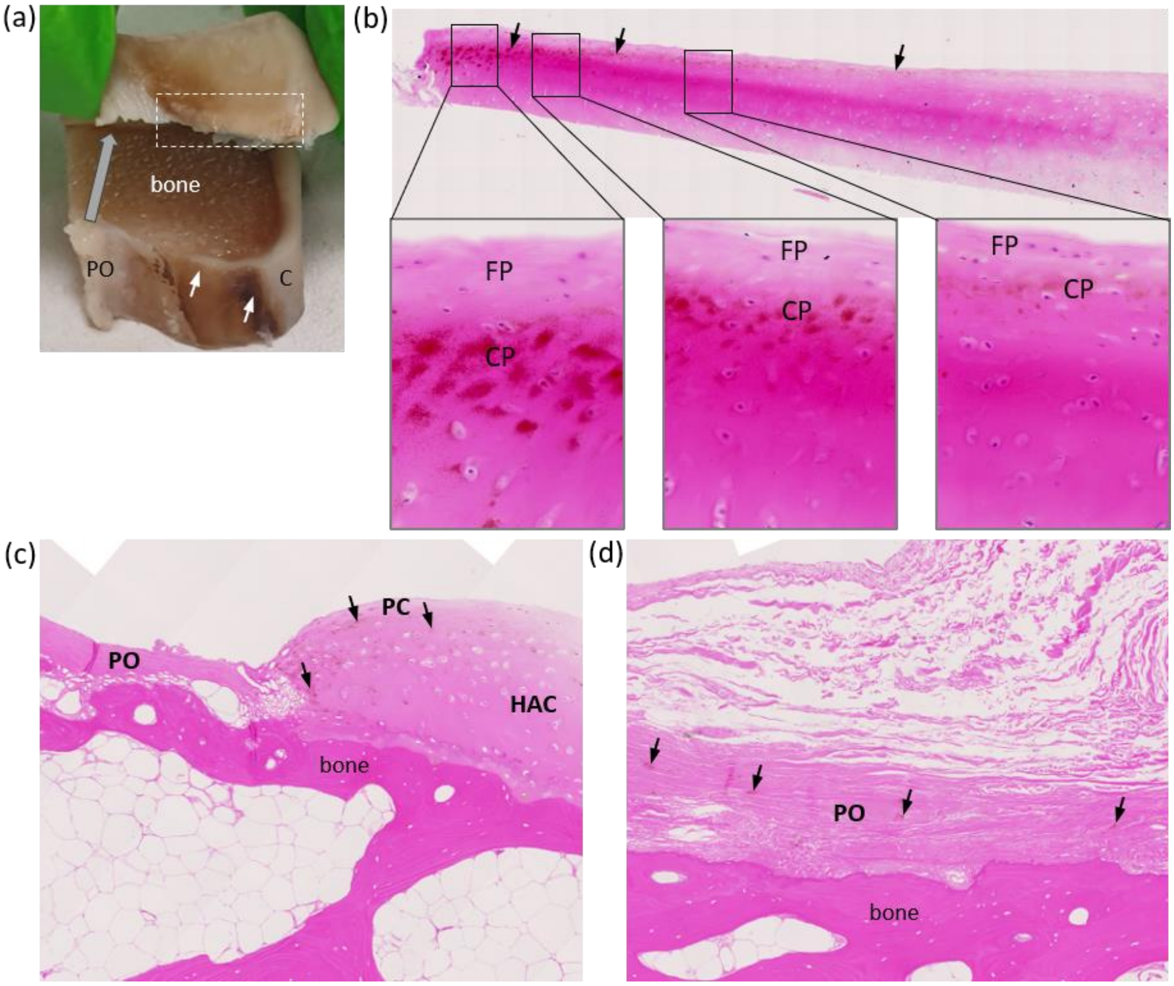


**Supplementary figure 5. Pigmentation of radial head perichondrium and periosteum**. Part of the radial head (post-fixation) is shown in (a), where it has been sliced and then a segment lifted up to show the cross section through the articular cartilage. The articular cartilage (C) appears white on the surface, until it reaches the periphery of the articular surface where it shows superficial pigmentation (white arrows). The more distal periosteum (PO) appears whiter. The top segment of the radial head (dashed box) was sectioned and stained with H&E, and is shown in (b), with the main articular surface towards the right of the image. Arrows indicate pigmentation of the perichondrium at the superficial edge of the cartilage which was lessens towards the articular surface and is no longer identifiable in the cartilage on the far right. The pigmentation here appears to be in the inner chondrogenic perichondrium (CP), rather than the outer fibrous perichondrium (FP). The lower segment of the radial head (not lifted up) in part (a) was sectioned and stained with H&E; (c) shows the junction of the perichondrium (PC) with the periosteum (PO), and (d) shows periosteum. Diffuse, granular pigmentation is observed in (c) within the perichondrium and adjacent hyaline articular cartilage (HAC). Small areas of pigmentation are observed within the periosteum, shown in (d).

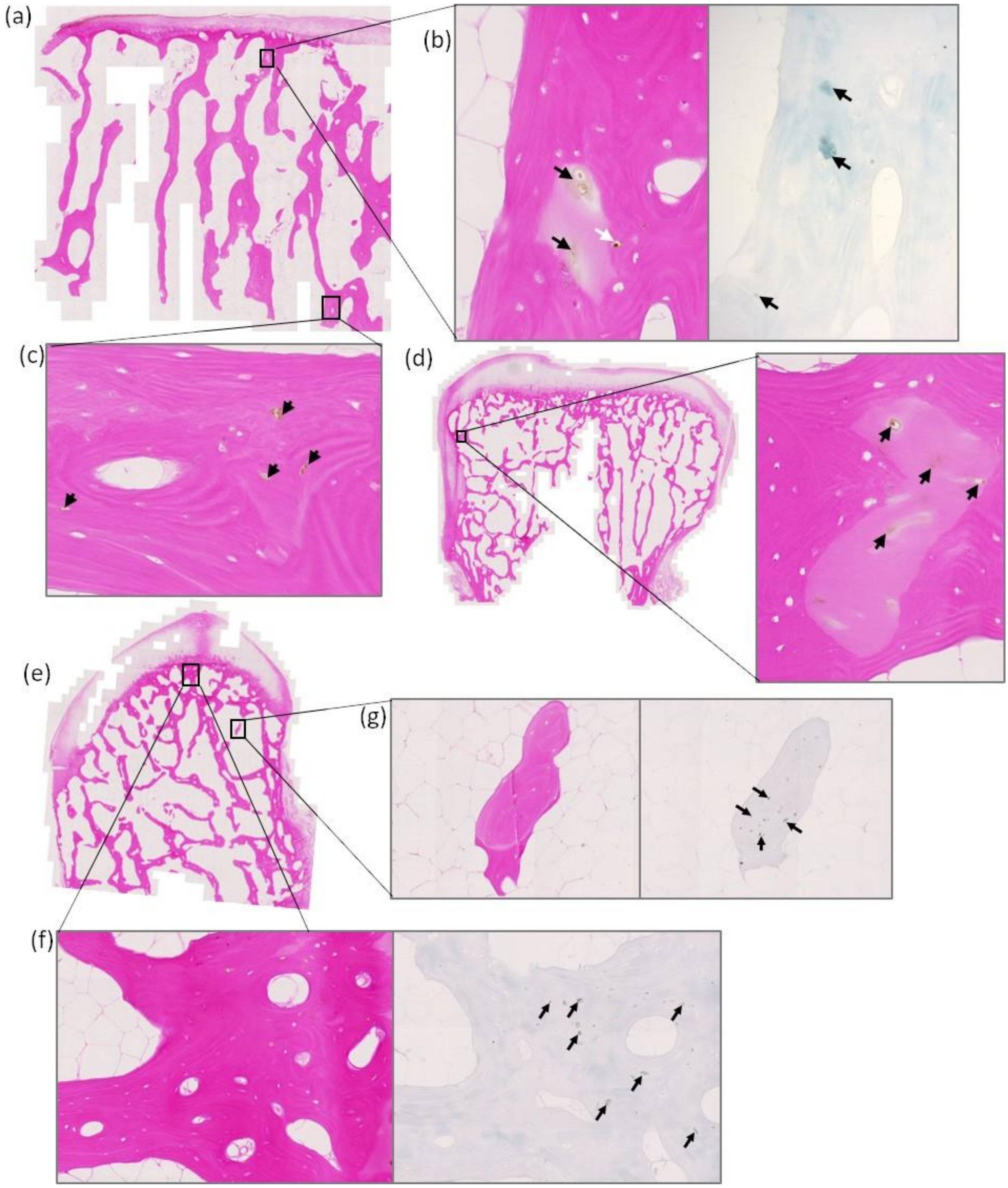


**Supplementary figure 6. Pigmentation of trabecular bone cells**. H&E staining of the lateral cuneiform is shown in (a). The insets from (a) show pigmented cells from the trabecular bone regions of the lateral cuneiform (a), with (b) showing chondrocytes that are not surrounded by eosinophilic lamellar bone and (c) showing osteocytes surrounding by mineralised lamellar bone. Schmorl's staining of the area in (b) is also shown. Pigmented chondrocytes within the radial head trabecular bone were also observed in (d), where the eosinophilic lamellar bone is not present and the cells appear large and round. H&E staining of the capitate is shown in (e) with (g) and (f) showing higher magnification images of trabecular bones regions with pigmented osteocytes, with corresponding Schmorl's staining of the same areas.

#### 1.4 Histology of viscera

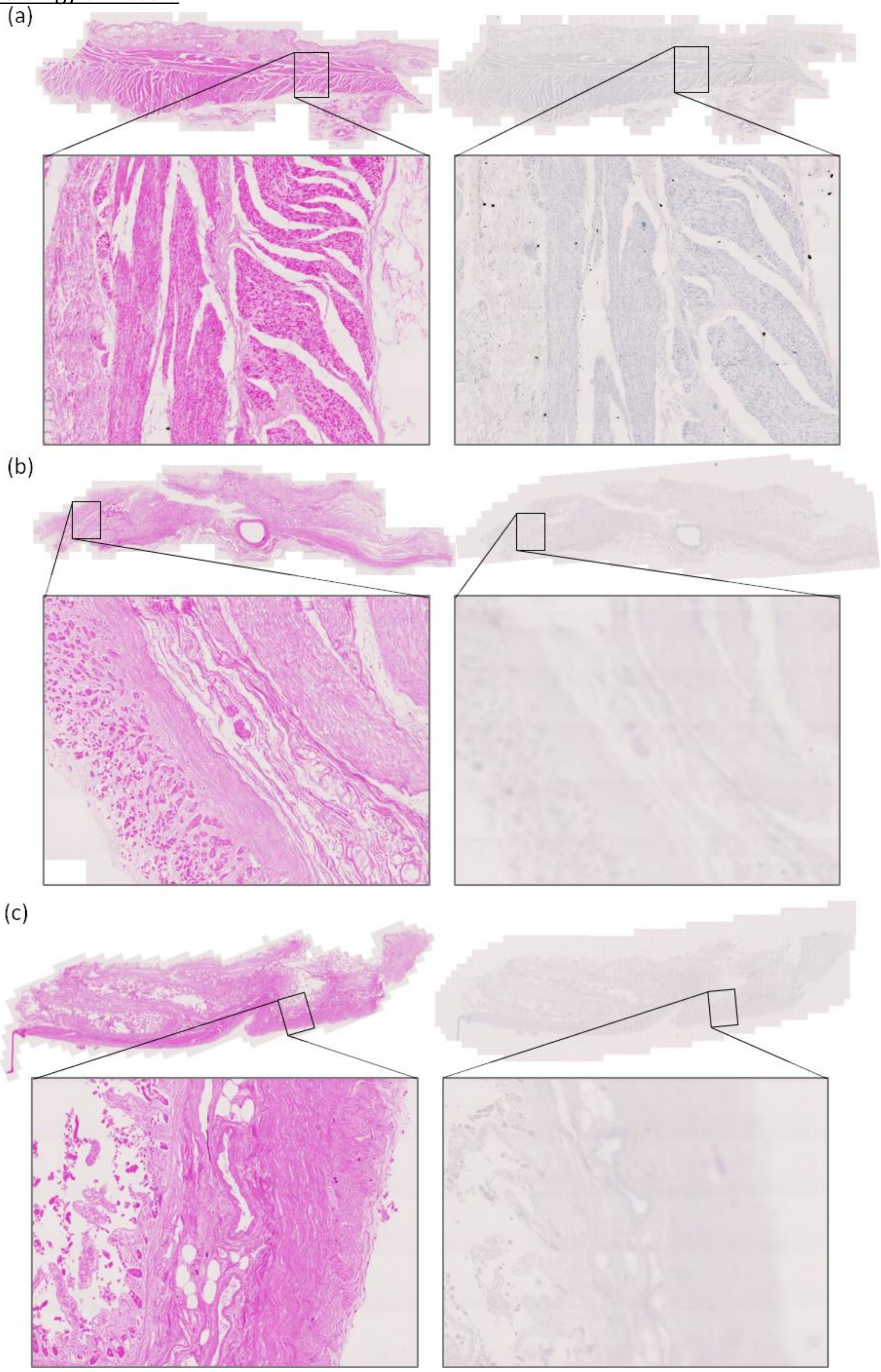


**Supplementary figure 7. Histology of the oesophagus, stomach and duodenum.** Haematoxylin & eosin (left column) and Schmorl's (right column) staining of the oesophagus (a), stomach (b), and duodenum (c). No ochronotic pigmentation was observed.

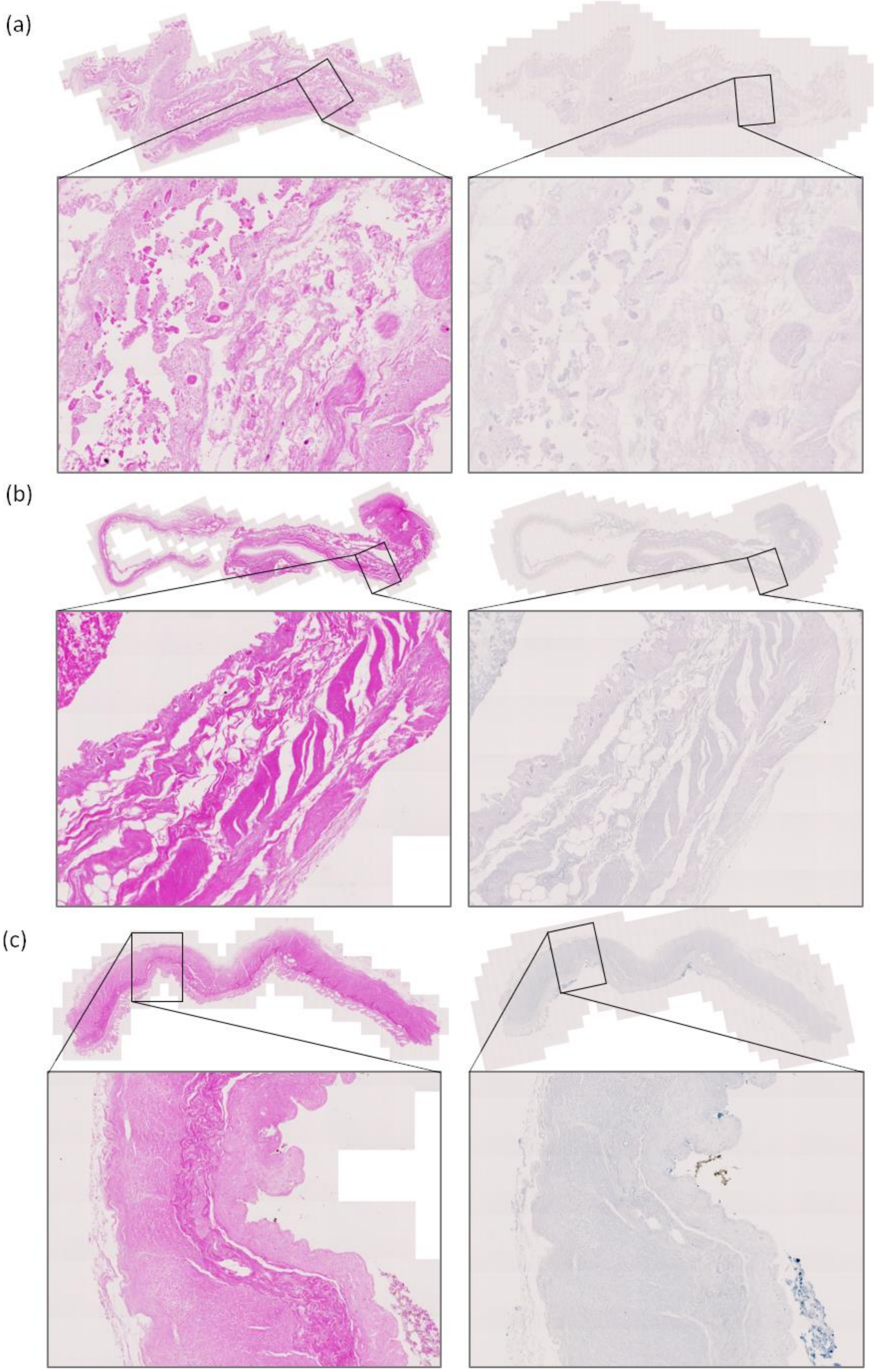


**Supplementary figure 8. Histology of the intestine.** Haematoxylin & eosin (left column) and Schmorl's (right column) staining of the jejunum (a), ileum (b) and large intestine (c), specifically the descending colon. No ochronotic pigmentation was observed.

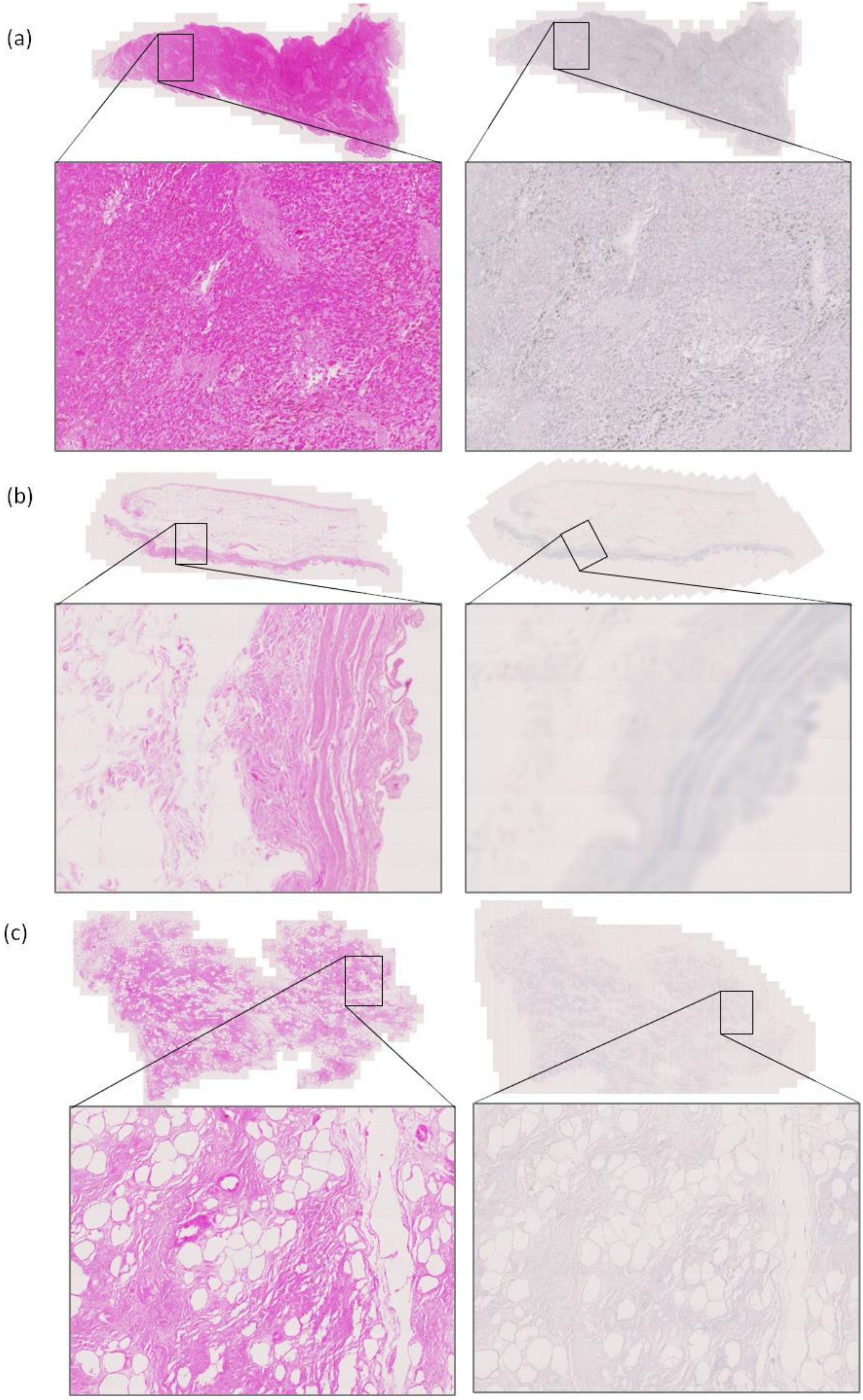


**Supplementary figure 9. Histology of hepatobiliary viscera.** Haematoxylin & eosin (left column) and Schmorl's (right column) staining of the liver (a), gallbladder (b) and pancreas (c). No ochronotic pigmentation was observed. The poor morphology of the liver suggests that necrosis had already set in by the point of biopsy of this tissue; the darker staining is not thought to be pigment, but signs of necrosis.

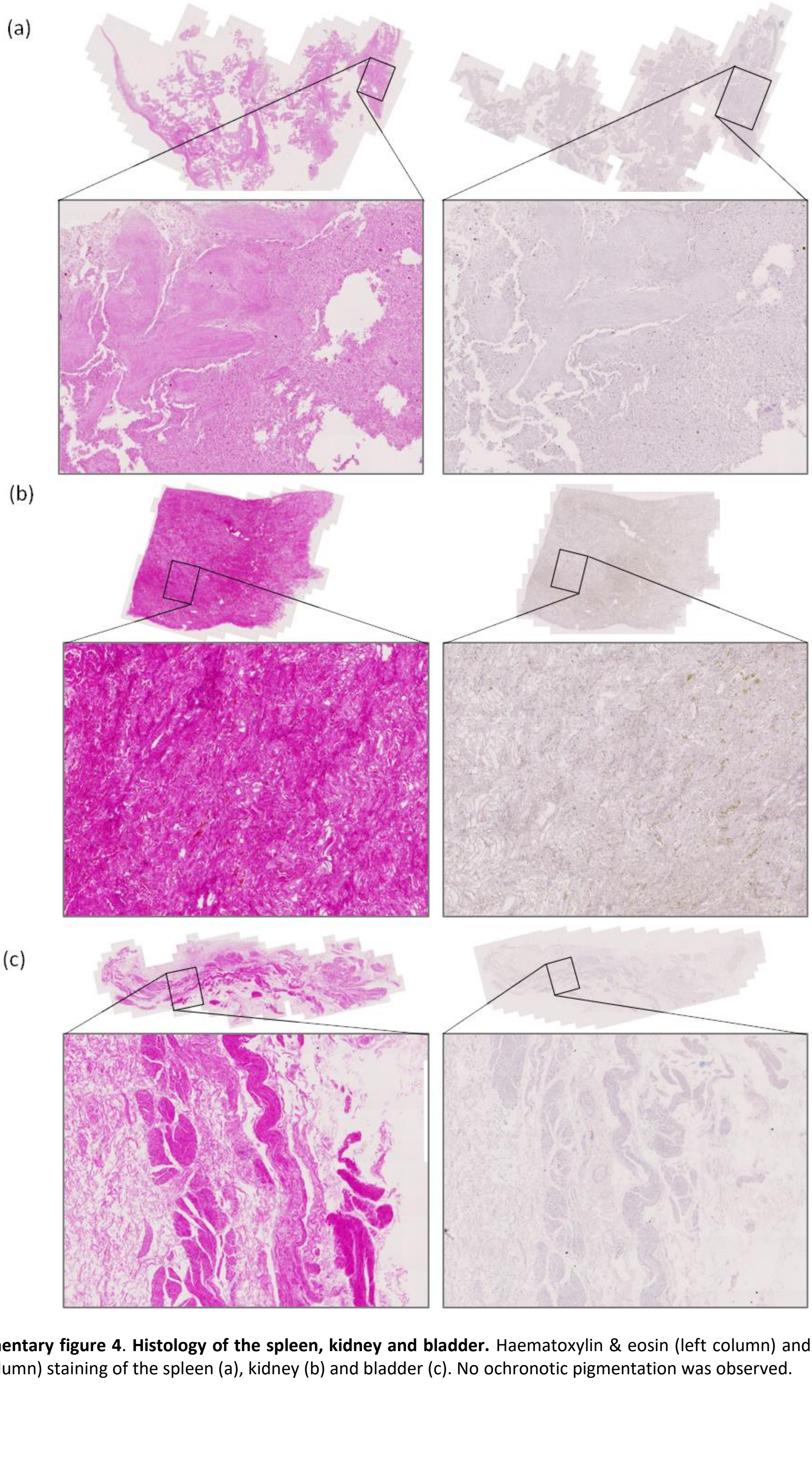


**Supplementary figure 4**. **Histology of the spleen, kidney and bladder.** Haematoxylin & eosin (left column) and Schmorl's (right column) staining of the spleen (a), kidney (b) and bladder (c). No ochronotic pigmentation was observed.

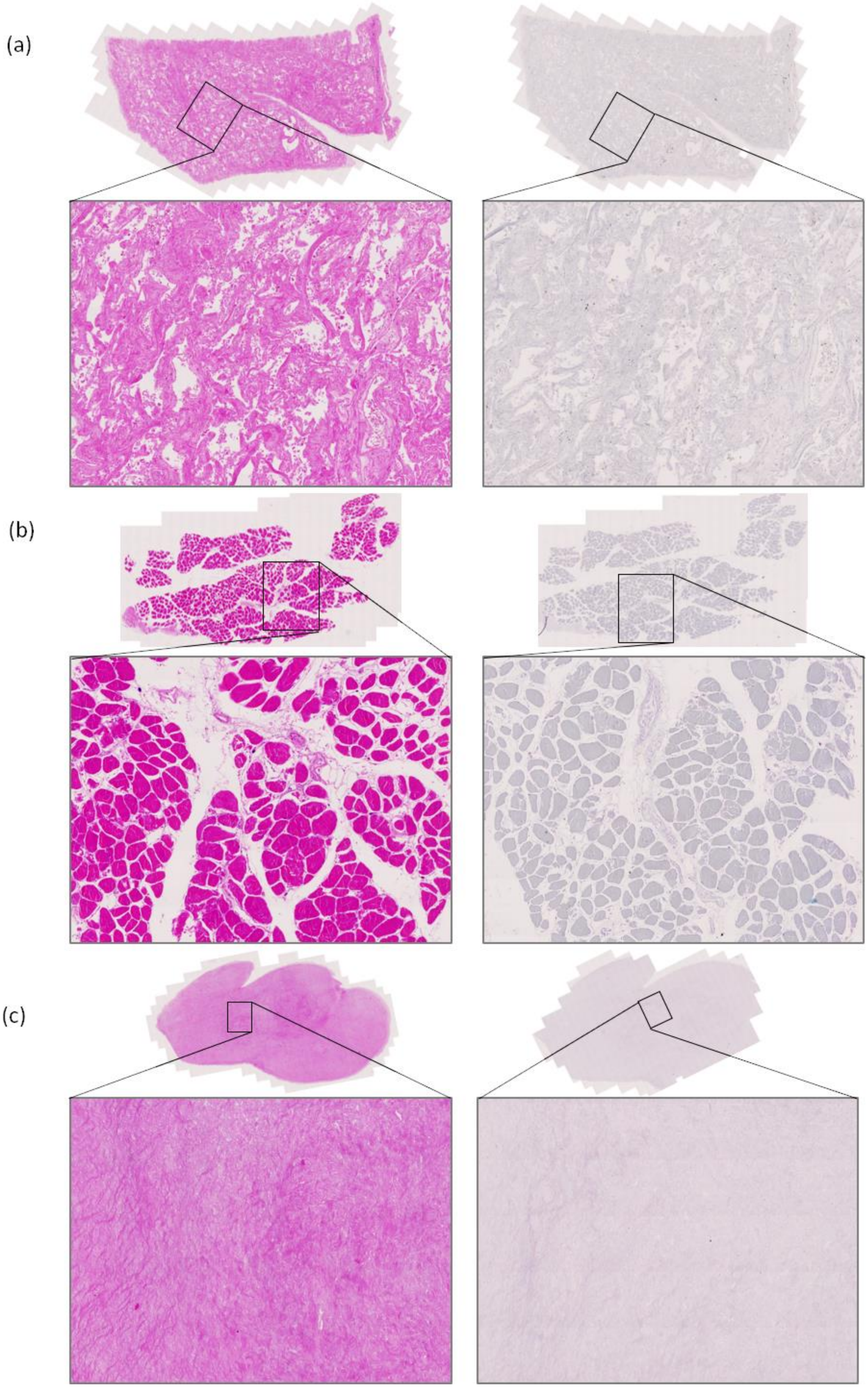


**Supplementary figure 11. Histology of lung, muscle and ovary.** Haematoxylin & eosin (left column) and Schmorl's (right column) staining of the lung parenchyma (a), skeletal muscle from the gastrocnemius near the Achilles tendon (b) and ovary (c). No ochronotic pigmentation was observed.

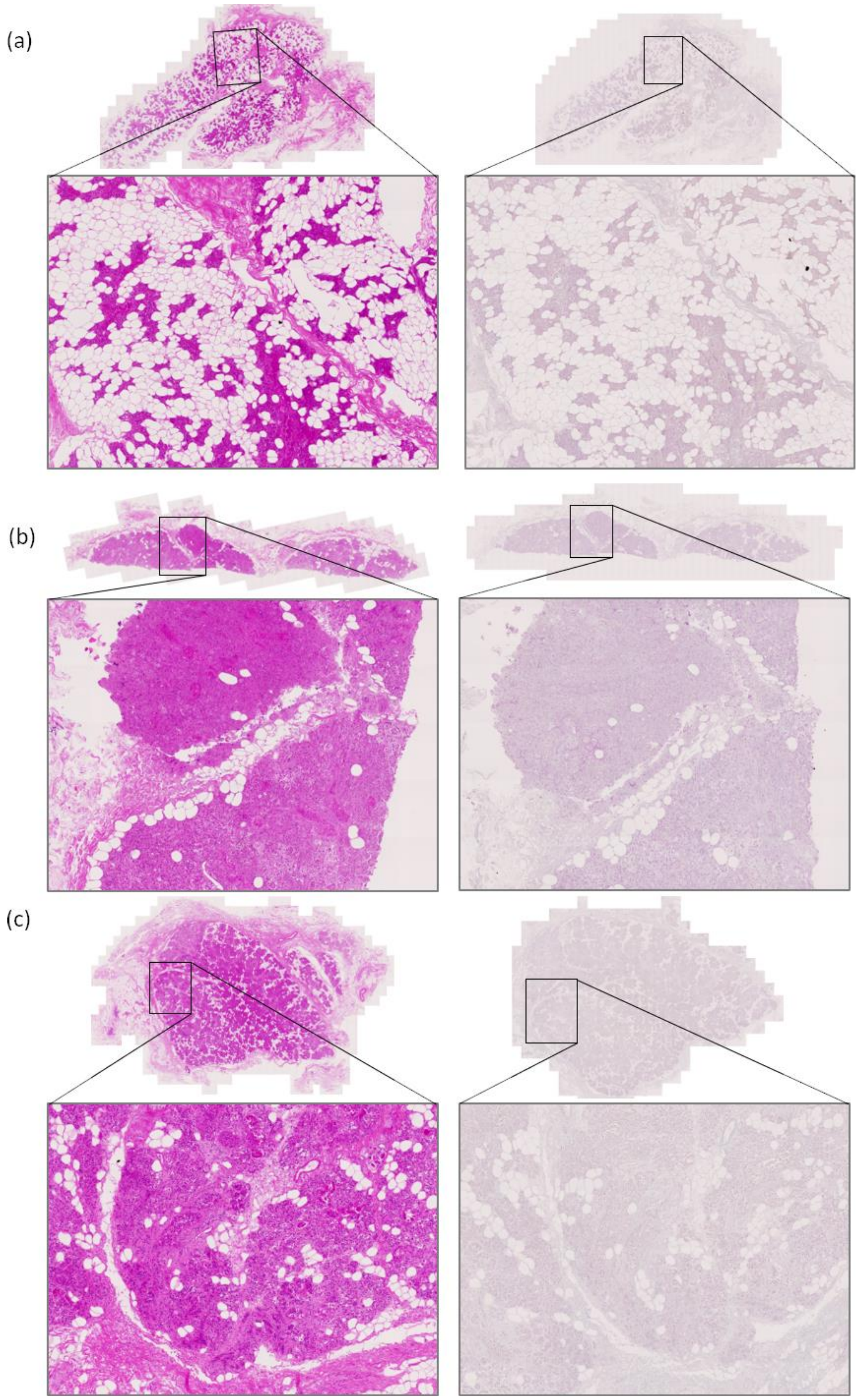


**Supplementary figure 12. Histology of salivary glands.** Haematoxylin & eosin (left column) and Schmorl's (right column) staining of the parotid gland (a), submandibular gland (b) and sublingual gland (c). No ochronotic pigmentation was observed.

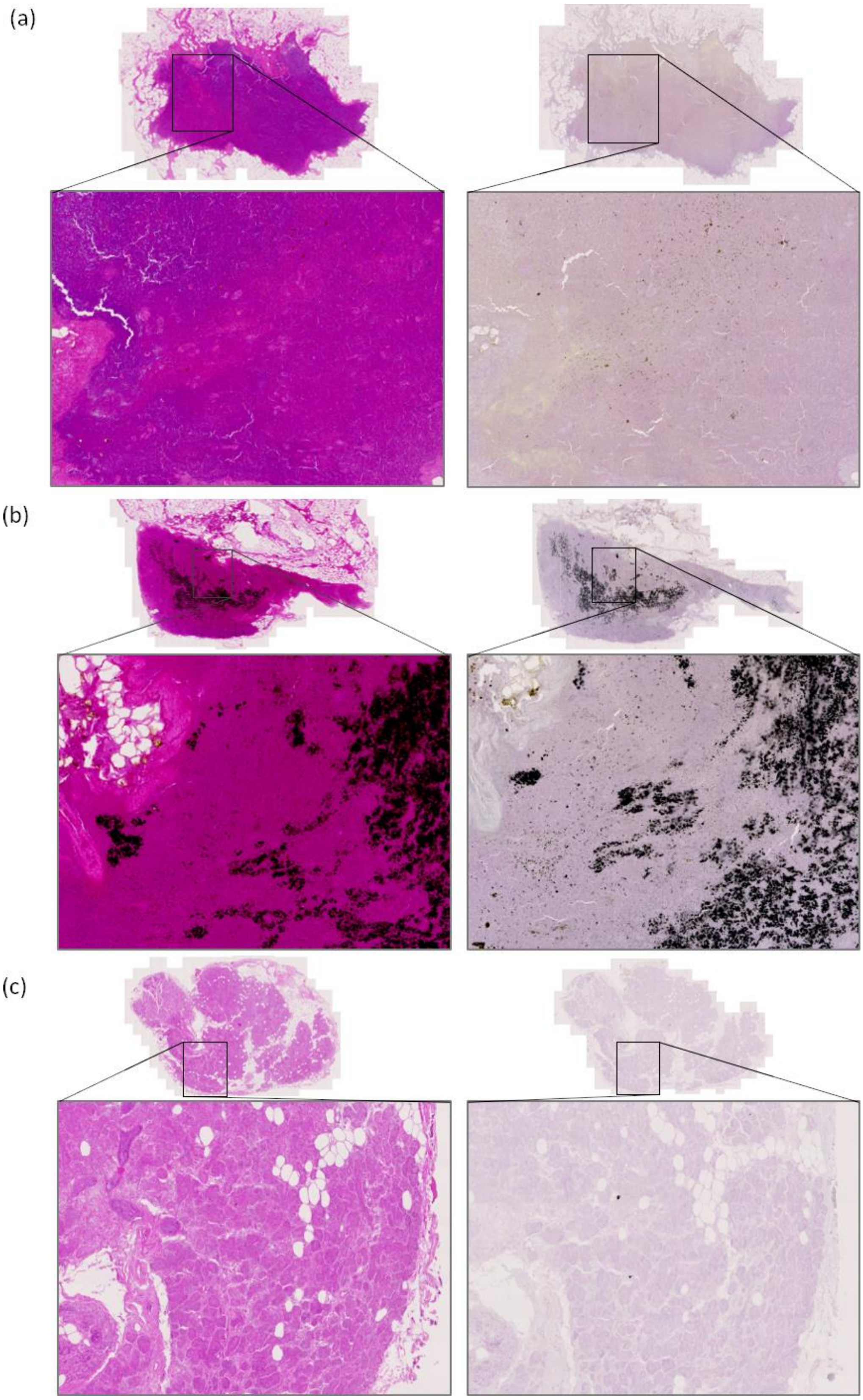


**Supplementary figure 13. Histology of lymph nodes and lacrimal gland.** Haematoxylin & eosin (left column) and Schmorl's (right column) staining of a carotid sheath lymph node (a), a tracheal lymph node (b) and the lacrimal gland (c). No ochronotic pigmentation was observed. The carotid lymph node had a yellow tinge, however the Schmorl's staining was purple in colour and not blue-green, indicating that it is not ochronosis. The tracheal lymph node had very intense black deposits, which are most likely carbon-based particle deposits, and not ochronosis.
